# Transforming Histology into Virtual Multiplex Immunofluorescence to Decode Prognostic Spatial Immunity in Hepatocellular Carcinoma

**DOI:** 10.64898/2026.02.25.707931

**Authors:** Linghan Cai, Songhan Jiang, Junhao Liang, Fengchun Liu, Buyi Zhang, Nic Gabriel Reitsam, Qinghe Zeng, Zheqi Hu, Yanqing Ma, Ziqian Li, Shi Feng, Maotong Hu, Xiuming Zhang, Jing Zhang, Jakob Nikolas Kather, Yongbing Zhang, Wenjie Liang

## Abstract

The spatial organization of the tumor immune microenvironment (TIME) drives hepatocellular carcinoma (HCC) prognosis but remains unquantifiable on routine H&E slides. Here, we present HCCExplorer, a deep learning framework that translates H&E into virtual multiplex immunofluorescence (mIF) and uses multi-modal graph learning to decode spatial survival signals. Trained on 30 H&E-mIF slide pairs, HCCExplorer evaluated a 1,813-slide multi-center cohort. It achieved superior overall survival stratification over clinical indices and state-of-the-art pathology and protein foundation models, yielding a concordance index of 0.71 and a Hazard Ratio (HR) of 15.46 (*P* < 0.001), maintaining stability across three external cohorts. Beyond risk stratification, interpretation of model features identified M1 macrophage infiltration as a protective determinant (HR=0.40, *P* < 0.05). Furthermore, it uncovered a protective “Containment Niche” at the invasion frontier (HR=0.02, *P* < 0.01), featuring macrophages co-localizing with Foxp3+ Tregs and CD4+ T cells. Ultimately, HCCExplorer provides actionable, spatially-resolved biomarkers from conventional histology for precision HCC management.

## Introduction

Hepatocellular carcinoma (HCC) is a leading cause of cancer mortality worldwide and remains a significant challenge for precision oncology owing to marked inter- and intra-tumoral heterogeneity and frequent underlying liver disease that constrains therapy choices^1–5^. Although immune checkpoint inhibitors (ICIs) have reshaped systemic treatment paradigms, only a subset of patients (< 30%) derive durable benefit^6^. This underscores the need for biomarkers that capture the biological determinants of response and progression^7–9^. A growing body of evidence indicates that these determinants are not solely molecular bulk readouts or average cell-type abundances, but are encoded in the spatial organization of the tumor immune microenvironment (TIME), including immune exclusion^10^, immune deserts^11^, and immunosuppressive myeloid niches^12^, which collectively influence tumor evolution^13^ and therapeutic resistance^14^. Therefore, decoding spatially resolved immunophenotypes in HCC is essential for robust risk stratification and for identifying clinically actionable and mechanistically interpretable biomarkers.

Spatial profiling technologies such as multiplex immunofluorescence (mIF) can quantify cell states while preserving tissue architecture, thereby enabling direct interrogation of interactions between immune and tumor cells^15–17^. However, their routine clinical translation is hindered by substantial costs, limited throughput, and operational complexity^18^. These factors restrict their availability and impede large-scale retrospective validation across diverse real-world cohorts. In contrast, haematoxylin and eosin (H&E) histopathology is ubiquitous, affordable, and deeply integrated into clinical workflows^19^. However, current H&E-based pathologic reporting faces critical bottlenecks: such assessment is typically qualitative; it neither quantifies immune infiltration levels nor captures the complex spatial interactions between tumor and immune cells. Furthermore, morphological patterns alone cannot resolve functionally distinct immune subsets. For example, it is impossible to distinguish immunosuppressive regulatory T cells from other lymphoid populations based solely on morphology^20^. These limitations obscure immune mechanisms that are central to prognosis and treatment response.

Recent breakthroughs in computational pathology indicate a transformative path for precision oncology by using multi-modal artificial intelligence (AI) to learn cross-modal mappings between morphology and molecular readouts^21–23^. VirtualMultiplexer^24^ employs contrastive unpaired translation to synthesize multiplexed immunohistochemistry (mIHC) images, accelerating histopathology workflows. GigaTIME^25^ demonstrated that deep learning models trained on massive paired H&E and mIF datasets can translate routine histology into virtual mIF across multiple protein channels, enabling population-scale analysis of the TIME. Complementarily, HEX^26^ showed that a foundation model (FM)-based pipeline can infer virtual spatial proteomics from H&E images, demonstrating that fusing histology with AI-derived spatial biomarkers improves prognostic accuracy, especially in lung cancer. While these studies collectively establish that routine histology encodes latent information predictive of spatial immune programs, translating virtual immunophenotyping into robust biomarkers for HCC presents distinct challenges. First, FM approaches often rely on extensive, pixel-aligned paired training data; however, curating such datasets at scale in clinical practice is constrained by inherent variability in tissue processing and registration complexities^27^. Second, moving beyond patch-level aggregates to capture granular cell-level interactions remains a critical frontier for maximizing the mechanistic interpretability of spatial biomarkers^28^. These challenges motivate the development of frameworks that are simultaneously data-efficient, topologically consistent, and explicitly spatial in modeling tumor-immune interplay.

Here, we present HCCExplorer, a spatially resolved and interpretable framework for virtual immunophenotyping from standard H&E slides and for the discovery of prognostic biomarkers in HCC. HCCExplorer introduces two key methodological advances. First, we develop a Cell-Consistent Cross-Modal Unpaired Translation (C3UT) method, which transforms H&E images into high-fidelity virtual mIF channels (e.g., CD3, CD4, and CD8). By enforcing a cell-map-guided consistency constraint, C3UT mitigates the risk of hallucination in virtual staining even with limited training data, enabling the characterization of a massive landscape of approximately 9 billion cells across 1,891 slides using 154,916 coarsely aligned patch pairs from 30 H&E-mIF whole-slide images. This generative ability bridges a methodological gap: while pathology FMs (e.g., UNI-h2^29^, CHIEF^30^, Prov-GigaPath^31^, and TITAN^32^) are confined to morphological features, and proteomics FMs (e.g., KRONOS^33^) necessitate physical antibody staining, C3UT uniquely infers molecular signals directly from standard histology. Second, we propose a multi-modal graph-based survival prediction model that captures spatial interactions among tumor and immune compartments, transforming static histology into prognostically informative interaction graphs.

We validated HCCExplorer on a multi-center cohort of 1,813 patients across four institutions. The framework consistently outperformed clinical standards and state-of-the-art pathology and protein FMs in prognostic stratification. Beyond prediction, decoding immune features revealed that macrophage infiltration is a robust protective determinant, characterized as M1 polarization through integrative analysis of bulk RNA sequencing data. Spatial analysis based on graph learning revealed a protective macrophage-orchestrated “Containment Niche” at the invasion frontier. By extracting these complex spatial signatures directly from routine histology, HCCExplorer establishes a scalable paradigm for precision oncology that bridges the gap between morphological patterns and molecular phenotypes in hepatocellular carcinoma.

## Results

### 1 Overview of HCCExplorer workflow

To understand the complex immune landscape of HCC and identify robust prognostic biomarkers from routine histology, we developed HCCExplorer, a multi-stage deep learning framework (Figure 1). This framework integrates generative AI with contextual multi-modal survival analysis to bridge the gap between morphological patterns and molecular profiles (Figure 1a-c).

**Figure 1.**
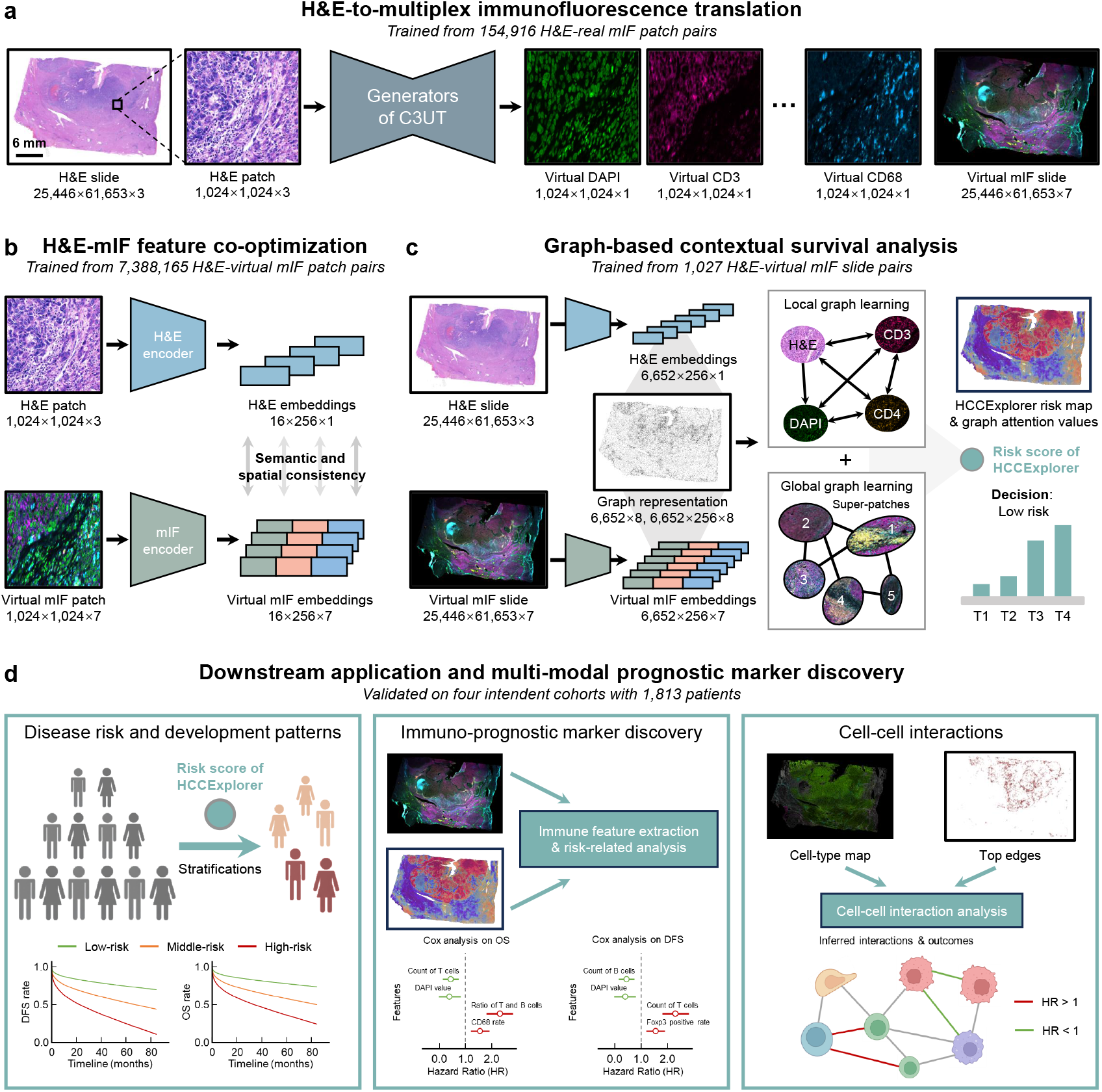
Overview of the HCCExplorer framework for virtual immunophenotyping and prognostic biomarker discovery. **a** H&E-to-multiplex immunofluorescence (mIF) translation. The framework employs a Cell-Consistent Cross-modal Unpaired Translation (C3UT) model to act as a virtual stainer. Trained on 154,916 real H&E-mIF patch pairs, the generator transforms standard H&E patches (1024 × 1024 × 3) into high-fidelity virtual mIF patches (1024 × 1024 × 7)), reconstructing the spatial expression of multiple immune markers (e.g., DAPI, CD3, and CD68) without physical staining. **b** H&E-mIF features co-optimization. To bridge morphological patterns with molecular profiles, feature encoders are pre-trained on a massive dataset of 7,388,165 H&E-virtual mIF patch pairs, each patch with 256 × 256 pixels. A contrastive learning strategy aligns the feature spaces of the two modalities, enforcing semantic and spatial consistency in the extracted embeddings (16 × 256 × 1for H&E and 16 × 256 × 7 for mIF, where 16 indicates the number of patches for an 1024 × 1024 large patch, 256is the feature dimension, and 7 is the number of mIF channels). **c** Graph-based contextual survival analysis. Multi-modal features are integrated via a hierarchical graph learning module trained on 1,027 slide pairs. Local graph learning captures H&E-virtual mIF relations in 256 × 256 patches, while global graph learning aggregates super-patches to model macro-level tissue architecture. The network outputs a comprehensive risk score and generates an interpretable spatial risk map for patient stratification. **d** Downstream application and multi-modal prognostic marker discovery. The HCCExplorer is validated on four independent cohorts comprising 1,813 patients. The framework provides interpretability through three avenues: (left) stratifying patients into low-, middle-, and high-risk groups with distinct survival outcomes; (middle) discovering immuno-prognostic markers via immune feature extraction and Cox regression analysis; and (right) inferring critical cell-cell interactions and their hazard ratios (HR) to elucidate the biological drivers of HCC progression.

The workflow begins with Cell-Consistent Cross-modal Unpaired Translation (C3UT), a H&E-to-virtual mIF staining module designed to recover latent molecular signals from routine histology (Figure 1a). Trained on 154,916 real H&E-mIF coarsely aligned patch pairs from 30 patients (Extended Data Figure 1a, Methods “H&E-to-virtual mIF translation”), C3UT transforms standard H&E patches into high-fidelity virtual mIF images. These inferred patches are spatially reassembled into whole-slide images (WSIs), enabling the precise mapping of key immune markers across the entire tumor landscape without requiring physical antibody staining.

Following virtual mIF translation, the second stage involves co-optimizing H&E and virtual mIF features to train robust feature encoders (Figure 1b). We constructed a large-scale pre-training dataset comprising 7,388,165 pixel-aligned H&E-virtual mIF patch pairs. By employing H&E-virtual mIF contrastive learning (Extended Data Figure 2a-c, Methods “H&E-virtual mIF contrastive learning”), we aligned the feature spaces of histological morphology and molecular phenotypes (Figure 1b). This co-optimization enforces semantic and spatial consistency, ensuring that the extracted features encapsulate both structural details and inferred immunophenotypes.

Building upon these aligned features, we implemented a graph-based multi-modal contextual survival prediction module (Figure 1c). Trained on 1,027 H&E-virtual mIF slide pairs, this module constructs a hierarchical graph representation of the HCC tissue (Extended Data Figure 2d, Methods “Graph-based multi-modal contextual survival analyses”). At the local level, the module captures cellular interactions (e.g., between CD3+ T cells and CD4+ T cells); at the global level, it aggregates spatially adjacent patches into super-patches to model the tissue’s macro-architecture. By integrating morphological and virtual immune features, HCCExplorer generates a comprehensive risk score and a spatial risk map to stratify patients into distinct risk categories (Figure 1c, Methods “Graph-based multi-modal contextual survival analyses”).

Finally, the framework was deployed for downstream applications and to discover immuno-prognostic markers (Figure 1d). We validated HCCExplorer on four independent cohorts, demonstrating its robustness in stratifying overall survival (OS) and disease-free survival (DFS) (Table 1). Beyond risk prediction, HCCExplorer offers interpretability through three key avenues: (i) delineating disease development patterns across risk groups; (ii) discovering immuno-prognostic markers via hand-crafted feature extraction; and (iii) inferring critical cell-cell interactions (Figure 1d). This interpretable design facilitates the identification of specific cellular neighborhoods and interaction motifs that either promote or suppress tumor progression, thereby providing actionable biological insights into HCC.

**Table 1.** Clinical characteristics of patients used in this study.

| Center | FAZJU |  | SAZJU | TCGA | YWCH |
| --- | --- | --- | --- | --- | --- |
| Data split | Train | Test | Test | Test | Test |
| Number of cases | 949 | 237 | 211 | 342 | 74 |
| Median age (years) | 57 (28-89) | 55 (21-86) | 67 (26-91) | 61 (16-90) | 69 (28-93) |
| Male | 800 (84.3%) | 199 (83.9%) | 185 (87.7%) | 232 (67.8%) | 59 (79.7%) |
| Female | 149 (15.7%) | 38 (15.1%) | 26 (12.3%) | 110 (32.2%) | 15 (20.3%) |
| Single tumor | 731 (77%) | 187 (78.9%) | 178 (84.3%) | — | 60 (81.1%) |
| Multiple tumors | 218 (23%) | 50 (21.1%) | 31 (14.7%) | — | 14 (18.9%) |
| Stage I | 603 (63.5%) | 159 (67.1%) | 137 (64.9%) | 166 (48.6%) | 59 (79.7%) |
| Stage II | 243 (25.6%) | 67 (28.3%) | 11 (5.2%) | 89 (26.0%) | 9 (12.2%) |
| Stage III | 103 (10.8%) | 11 (4.6%) | 63 (29.9%) | 87 (25.4%) | 6 (8.1%) |
| Grade I | 65 (6.9%) | 19 (8.0%) | 31 (14.7%) | 165 (48.2%) | 22 (29.7%) |
| Grade II | 437 (46.0%) | 108 (45.6%) | 134 (63.5%) | 118 (34.5%) | 33 (44.6%) |
| Grade III | 368 (38.8%) | 90 (38.0%) | 29 (13.7%) | 46 (13.5%) | 4 (5.4%) |
| Grade IV | 79 (8.3%) | 20 (8.4%) | 17 (8.1%) | 13 (3.8%) | 15 (20.3%) |
| OS state |  |  |  |  |  |
| Censor | 186 (19.6%) | 49 (20.7%) | 52 (24.6%) | 119 (34.8%) | 49 (66.2%) |
| Median follow-up | 47 (0.5-122) | 47 (1-125) | 72 (3-114) | 19.74 (0-120.7) | 62 (2-110) |
| NA | 0 (0%) | 0 (0%) | 0 (0%) | 0 (0%) | 0 (0%) |
| DFS state |  |  |  |  |  |
| Censor | 575 (60.6%) | 146 (61.6%) | 117 (55.6%) | 163 (47.7%) | 42 (56.8%) |
| Median follow-up | 25 (0-122) | 27 (0-125) | 58 (0.65-162) | 14.7 (0-120.7) | 14.4 (1.57-64.5) |
| NA | 0 (0%) | 0 (0%) | 0 (0%) | 44 (12.7%) | 7 (9.5%) |

### 2 HCCExplorer transforms hepatocellular carcinoma pathology slides into virtual multiplex immunofluorescence slides

To reconstruct the tumor immune microenvironment (TIME) directly from standard histology, we trained the C3UT module of HCCExplorer using 30 coarsely aligned H&E-mIF pairs from the internal First Affiliated Hospital of Zhejiang University (FAZJU) cohort (Figure 1a, Table 1, Methods “H&E-to-virtual mIF translation”). The model was optimized to predict the expression of seven clinically relevant markers (Figure 1a). Visually, the virtual mIF images generated by HCCExplorer successfully recapitulated the tissue morphology of the original H&E slides while preserving staining patterns observed in real mIF references (Figure 2a, Extended Data Figure 3a).

**Figure 2.**
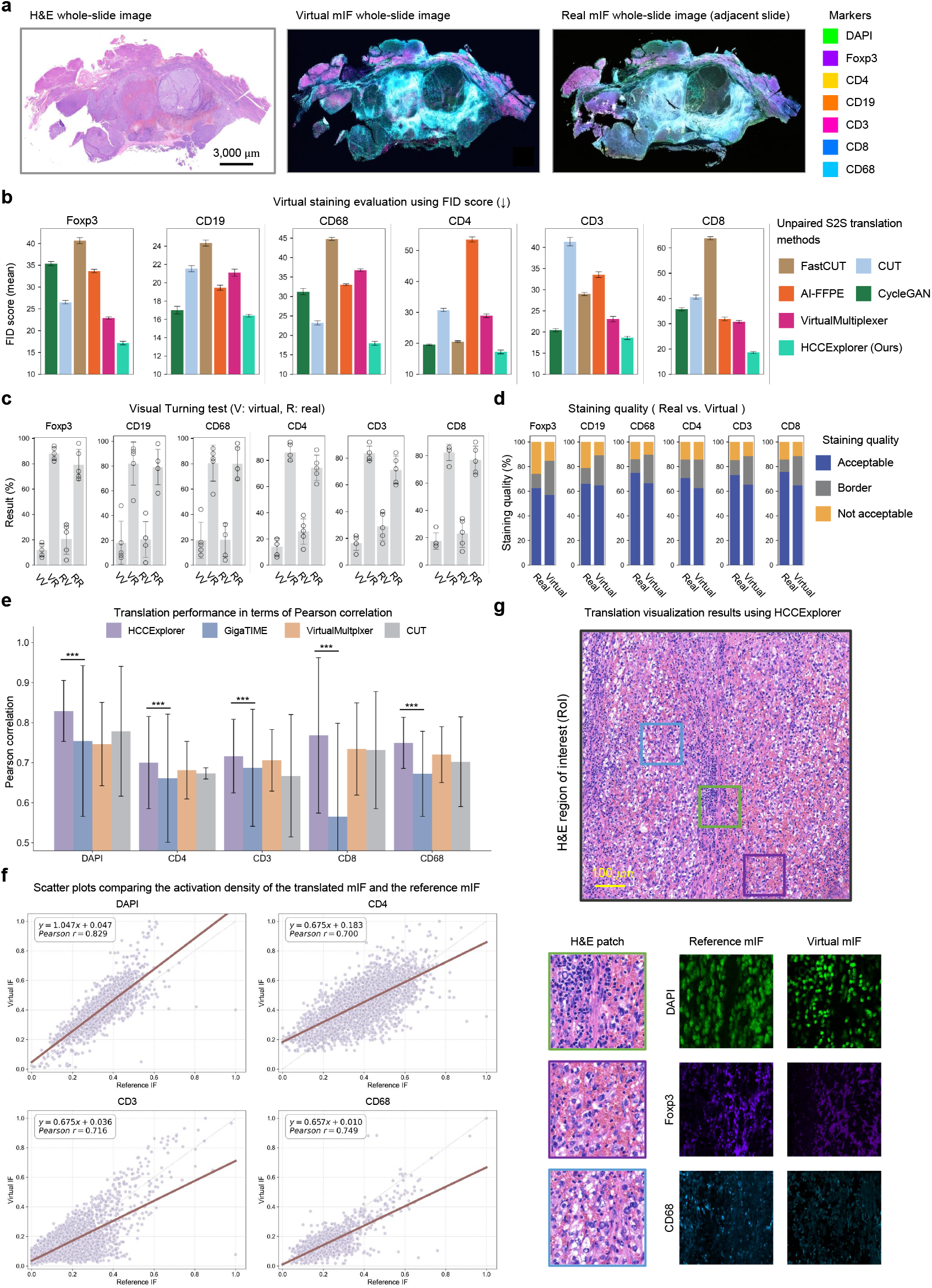
Performance evaluation and biological validation of HCCExplorer in transforming H&E images to virtual multiplex immunofluorescence. **a** Whole-slide visualization of virtual staining. Representative images display an original H&E slide (left), the HCCExplorer-generated virtual mIF slide (middle), and the corresponding real mIF slide from an adjacent tissue section (right). The model simultaneously reconstructs seven markers (DAPI, Foxp3, CD4, CD19, CD3, CD8, and CD68), color-coded as indicated in the legend, effectively recapitulating the tumor immune microenvironment structure. **b** Quantitative benchmarking of generation quality. Fréchet Inception Distance (FID) scores (lower indicates better quality) comparing HCCExplorer against five state-of-the-art unpaired image translation methods across six markers. HCCExplorer consistently achieves the lowest FID scores. Data are presented as mean ± standard deviation (std). **c** Visual Turing test results. Assessment by four expert pathologists showing the percentage of image patches classified as “Real” for Virtual (V) generated images versus Real (R) authentic images. Experts perceived a high proportion of virtual images as authentic, indicating no significant perceptual boundaries. **d** Staining quality assessment. Pathologists’ evaluation of staining quality (categorized as Acceptable, Borderline, or Not acceptable) comparing Real versus Virtual images based on expression levels and subcellular localization. Virtual staining quality is comparable to, and in some markers (e.g., CD8, CD68) surpasses, physical staining. **e** Pixel-level translation accuracy. Pearson correlation coefficients (PCC) comparing the translation performance of HCCExplorer against GigaTIME, VirtualMultiplexer, and CUT. HCCExplorer demonstrates significantly higher concordance across all markers (\*\*\**P* < 0.001, ** *P* < 0.01, * *P* < 0.05, Williams’s t-test). Error bars represent standard deviation. **f** Activation density correlation. Scatter plots comparing the normalized signal intensity of virtual mIF (y-axis) versus reference mIF (x-axis) for DAPI, CD4, CD3, and CD68. The Pearson correlation coefficient (*r*) and linear regression equations indicate strong linear correlation in expression prediction. **g** High-resolution local visualization. A representative H&E region of interest (top) with magnified insets (bottom) showing paired H&E patches, reference mIF, and virtual mIF for specific markers (DAPI, Foxp3, CD68). The virtual images accurately capture fine-grained cellular details and morphology consistent with the ground truth.

To quantitatively evaluate translation quality, we conducted a comprehensive benchmark against five state-of-the-art unpaired image-to-image translation methods (FastCUT^34^, CUT^34^, AI-FFPE^35^, CycleGAN^36^, and VirtualMultiplexer^24^) and the recently published H&E-mIF paired translation model, GigaTIME^25^ (Figure 2c, Extended Data Figure 3b, c, Methods “Evaluation metrics”). In terms of distribution distance, HCCExplorer consistently achieved the lowest Fréchet Inception Distance^37^ (FID) scores across all markers compared to the unpaired image-to-image translation baselines, indicating the closest distributional alignment (Figure 2b). Notably, although GigaTIME benefits from paired data supervision, HCCExplorer demonstrated superior performance on overlapping markers in the HCC cohort (Extended Data Figure 3). For instance, HCCExplorer achieved a mean FID of 17.39 compared to 39.60 for GigaTIME (Extended Data Figure 3b). We further assessed structural preservation using the Contrast-Structure Similarity^38^ (CSS) index, where HCCExplorer surpassed competing models, including VirtualMultiplexer (e.g., CD68 CSS: HCCExplorer 79.81% vs. VirtualMultiplexer 70.15%), GigaTIME (e.g., CD68 CSS: HCCExplorer 79.81% vs. GigaTIME 66.78%), indicating that HCCExplorer yielded virtual images exhibiting greater structural fidelity to the ground truth in HCC (Extended Data Figure 3c).

To assess the clinical indistinguishability of the translated images, we conducted a visual Turing test (Figure 2c). Four pathologists with more than five years of specialized experience in liver cancer were presented with randomized real and virtual mIF patches, along with corresponding real H&E patches, and were tasked with classifying their origin (Methods “Visual Turing test”). As shown in Figure 2c, the experts’ assessment revealed no significant perceptual boundaries between the modalities. While experts correctly identified 76.80% of the real mIF images, they perceived 83.73% of the virtual images as real. The high acceptance rate corresponded to low sensitivity (16.27%) for detecting virtual artifacts, demonstrating that the virtual immunophenotypes generated by HCCExplorer possess textural and structural characteristics that are effectively indistinguishable from physical biological staining. Moreover, in a staining quality assessment (Figure 2d, Methods “Staining quality evaluation”), where pathologists evaluated labeled real and virtual images based on expression levels and subcellular localization, HCCExplorer demonstrated “Acceptable” quality rates comparable to those of real mIF slides, and even surpassing them in markers such as CD8 and CD68, which are often prone to artifacts.

Beyond visual similarity, we validated the biological fidelity of the translation at the pixel and cellular levels. We compared the translation performance using Pearson correlation coefficients (PCC) against advanced translation methods (Figure 2e). HCCExplorer yielded significantly higher correlation scores across all markers (*P* < 0.001; Figure 2e). Specifically, for DAPI (cellular staining), our model achieved high concordance, confirming its ability to precisely localize individual cells. To further verify that the model captures accurate expression levels rather than merely texture, we analyzed the activation density (Figure 2f). Scatter plots of signal intensity between virtual and reference mIF showed significant correlations (DAPI: *r* = 0.829 ± 0.076; CD68: *r* =0.749 ± 0.064; CD3: *r* =0.716 ± 0.092). These statistical metrics are corroborated by the model’s performance across various regions of interest (RoIs), in which the high-resolution translation accurately captures fine-grained cellular details consistent with the reference (Figure 2g). In summary, these results demonstrate that HCCExplorer learns generalizable, non-trivial relationships between tissue morphology and protein expression, enabling robust virtual phenotyping for downstream analyses (Figure 2e-g).

### 3 HCCExplorer identifies high-risk patients through H&E-virtual mIF multi-modal prognosis

To translate the spatially resolved virtual immunophenotypes into actionable clinical insights, we evaluated the prognostic efficacy of HCCExplorer in a multi-cohort study comprising 1,813 patients (Table 1). We developed a graph-based survival prediction model that integrates morphological embeddings from H&E images with co-optimized virtual mIF features (Figure 1b, c, Extended Data Figure 2d, Methods “Graph-based multi-modal contextual survival analyses”). The model was trained on the First Affiliated Hospital of Zhejiang University (FAZJU) cohort (949 patients) and validated on an internal test set (237 patients) as well as three independent external cohorts: Second Affiliated Hospital of Zhejiang University (SAZJU) (211 patients), The Cancer Genome Atlas Liver Hepatocellular Carcinoma (TCGA-LIHC) (342 patients), and Yiwu Central Hospital (YWCH) (74 patients) (Tables 1 and 2).

**Table 2.**
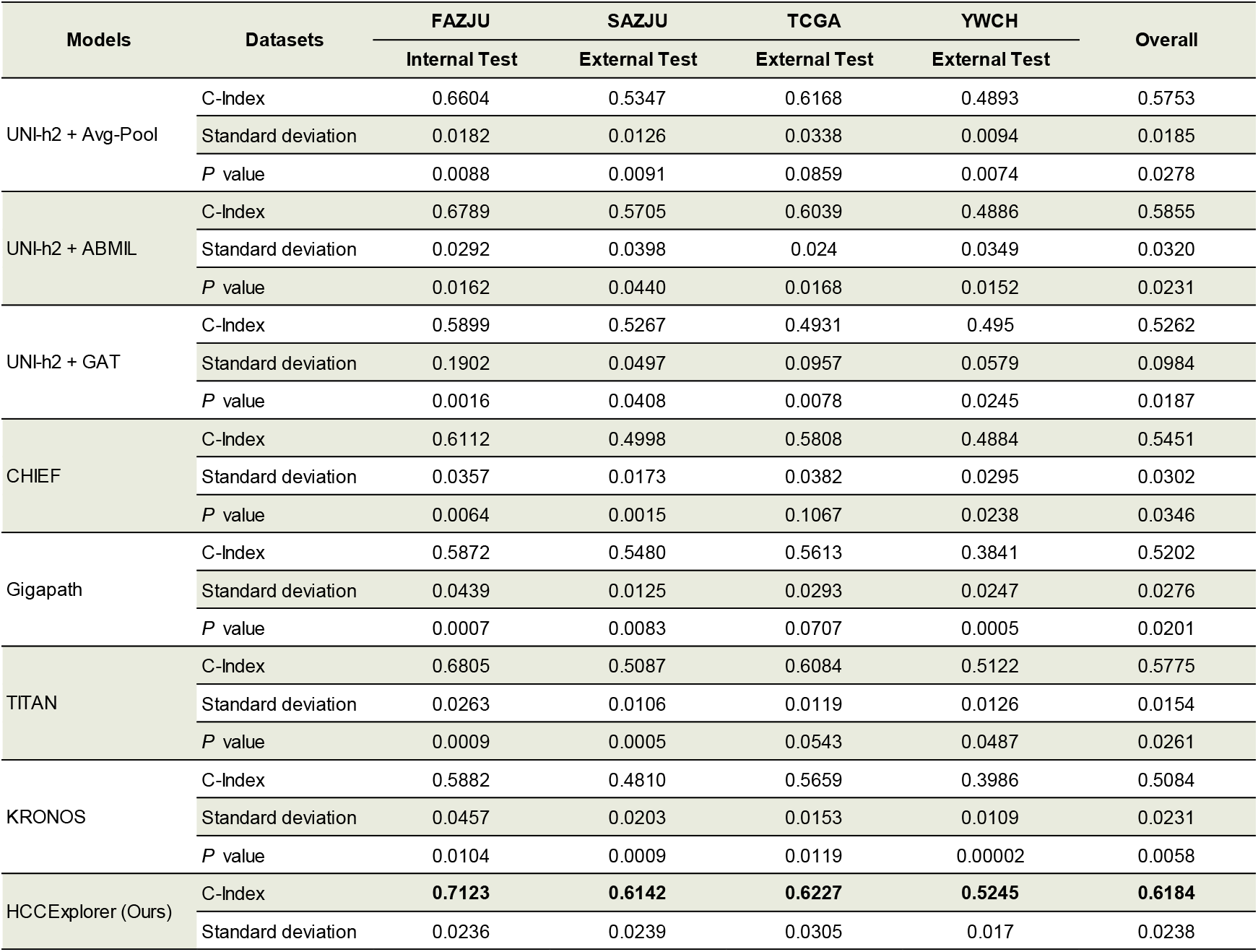
OS prediction comparison with other advanced methods. The models are trained on the FAZJU internal training set and directly tested on the test set of each cohort to evaluate generalization. *P* values were calculated through two-sided paired t-test.

We first evaluated whether HCCExplorer could improve risk stratification beyond standard clinical parameters (Figure 3a-c, Methods “Evaluation metrics”). In the FAZJU test set, HCCExplorer achieved a concordance index (C-index) exceeding 0.70 for overall survival (OS), disease-free survival (DFS), and recurrence-free survival (RFS), consistently outperforming standard histopathological variables, including AJCC stage^39^ and differentiation grade (Figure 3a). Notably, integrating risk scores from HCCExplorer with clinical variables (“All + HCCExplorer”) yielded the highest predictive performance (DFS: C-index=0.83, 95% confidence interval (CI): 0.74–0.90), suggesting that the model captures sub-visual prognostic signals in H&E-virtual mIF images (Figure 3a). In Cox regression analysis, the risk score of HCCExplorer demonstrated the strongest discriminatory power, achieving the highest Hazard Ratio of 15.46 (95% CI: 5.42–55.70, *P* < 0.001) for OS, 4.10 (95% CI: 1.77–9.58, *P* < 0.001) for DFS, and 4.22 (95% CI: 2.41–7.39, *P* < 0.001) for RFS among all tested covariates (Figure 3b).

**Figure 3.**
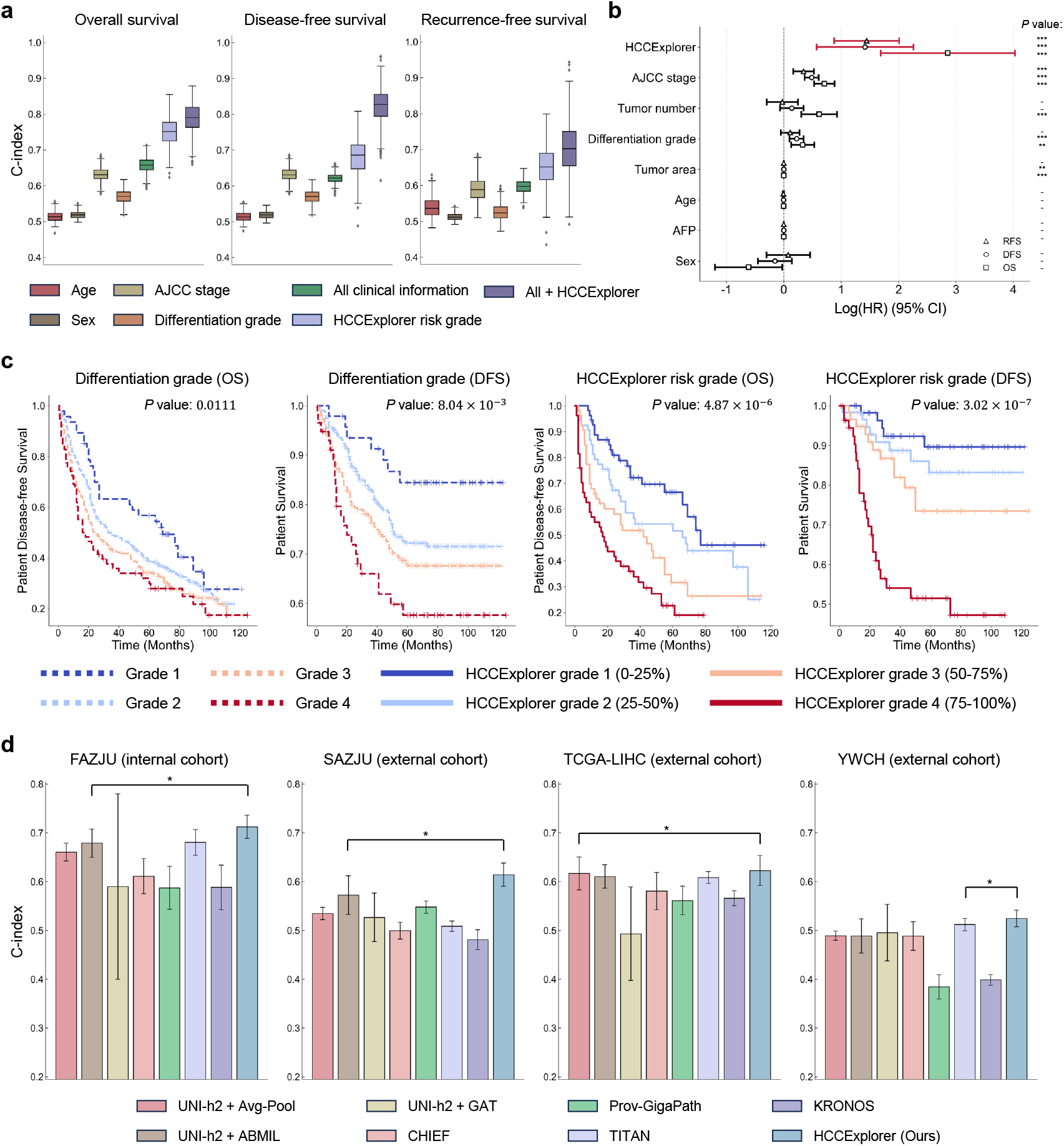
Prognostic performance and risk stratification capability of HCCExplorer in hepatocellular carcinoma. **a** Comparison of predictive accuracy using the concordance index (C-index). Boxplots illustrate the performance of standard clinical characteristics (age, sex, AJCC stage, differentiation grade), the HCCExplorer risk grade alone, and integrated models (“All + HCCExplorer”) across three survival outcomes: overall survival (OS), disease-free survival (DFS), and recurrence-free survival (RFS). The integrated model achieves the highest prognostic consistency, indicating that HCCExplorer captures sub-visual signals complementary to routine clinical variables. **b** Cox regression analysis of prognostic factors. The forest plot displays the Log hazard ratio (HR) with 95% confidence intervals (CI) for various covariates. The HCCExplorer risk score demonstrates the strongest discriminatory power among all tested variables, surpassing traditional markers such as AJCC stage and tumor number. Wald test^88^ was used to calculate *P* values for each covariate. **c** Kaplan-Meier survival analysis comparing histological grading versus HCCExplorer stratification. Left. Survival curves stratified by standard histological differentiation grade for OS and DFS show suboptimal outcomes (*P* =0.0111 and *P=* 8.04 × 10 ^−3^, respectively). Log-rank Test is used for *P* value calculation. Right. Survival curves stratified by HCCExplorer Risk Grade, where patients are discretized into four quartiles (Grade 1: 0-25% to Grade 4: 75-100%). HCCExplorer achieves distinct separation between risk groups with high statistical significance (*P =*4.87 × 10^−6^ for OS; *P*= 3.02 × 10^−7^ for DFS), effectively identifying high-risk patients prone to poor outcomes. **d** Survival prediction comparison of HCCExplorer and the state-of-the-art methods in four independent cohorts. The HCCExplorer obtains the best performance in the comparisons, surpassing the pathology foundation models (UNI-h2, CHIEF, Prov-GigaPath, and TITAN) and the protein foundation model (KRONOS). Two-sided paired t-test is used for *P* values calculation. Significance levels for this figure: * *P* < 0.05, ** *P* < 0.01, *** *P* < 0.001.

To assess clinical utility, we stratified patients into four risk quartiles based on HCCExplorer risk scores (Figure 3c). Kaplan-Meier (KM) analysis demonstrated that HCCExplorer yielded a more distinct separation of patient outcomes than histological differentiation grade (Figure 3c). While standard differentiation grading showed overlapping survival curves (particularly between Grades 1 and 2;), HCCExplorer achieved distinct stratification for both OS (*P* =4.87 × 10^−6^) and DFS (*P* = 3.02 × 10^−7^). These indicate that HCCExplorer offers greater prognostic granularity than conventional histological grading, making it a more reliable tool for risk assessment.

To validate the necessity of the proposed multi-modal strategy, we benchmarked HCCExplorer against seven state-of-the-art survival prediction methods, comprising multiple-instance learning (MIL)-based methods (ABMIL^40^, and GAT^41^), pathology foundation models (UNI-h2^29^, CHIEF^30^, Prov-GigaPath^31^, and TITAN^32^), and a proteomics foundation model KRONOS^33^ (Figure 3d, Table 2, Extended Data Figures 4, 5). HCCExplorer demonstrated excellent performance, achieving a C-index of 0.71 (95% CI: 0.66–0.74) in the internal cohort and maintaining the highest mean C-index of 0.58 across three external cohorts. Our framework outperformed pathology-based models (*P* < 0.05) such as TITAN (C-index: Internal = 0.68, 95% CI: 0.63–0.73; External = 0.54) and Prov-GigaPath (C-index: Internal = 0.59, 95% CI: 0.50–0.67; External = 0.50), which exhibited limited generalizability in multi-center settings (Figure 3d, Table 2, Extended Data Figures 4, 5). Crucially, the proteomics foundation model KRONOS also failed to surpass morphology-based baselines (C-index: Internal = 0.59, 95% CI: 0.50–0.68; External = 0.48) despite leveraging molecular signals (Figure 3d, Table 2). These results establish a critical biological insight: neither morphology alone (as in TITAN and Prov-GigaPath) nor molecular signals devoid of spatial context (as in KRONOS) are sufficient for precise prognostication. Only by explicitly modeling the latent immune landscape while retaining slide-level spatial information does HCCExplorer recover the multi-modal prognostic contexts necessary for robust survival analysis (Figure 3d, Table 2, Extended Data Figures 4, 5).

### 4 HCCExplorer captures immuno-prognostic features and identifies macrophage infiltration as a dominant biomarker

A major challenge in applying deep learning to clinical pathology is the “black box” nature of risk prediction^42^. To elucidate the biological rationale underlying HCCExplorer, we established an interpretability framework that links the model’s attention mechanism to quantifiable immunophenotypes (Figure 4a, Methods “Immune feature extraction” and “Immune feature analyses”). We hypothesized that the model assigns higher attention scores to tissue regions exhibiting specific characteristics of the TIME that are determinative of patient survival. To verify this, we extracted immune features from each patch with the highest and lowest attention scores, thereby translating abstract computational weights into tangible immune signatures (Figure 4a, Methods “Immune feature extraction” and “Immune feature analyses”, Supplementary Information Table S3).

**Figure 4.**
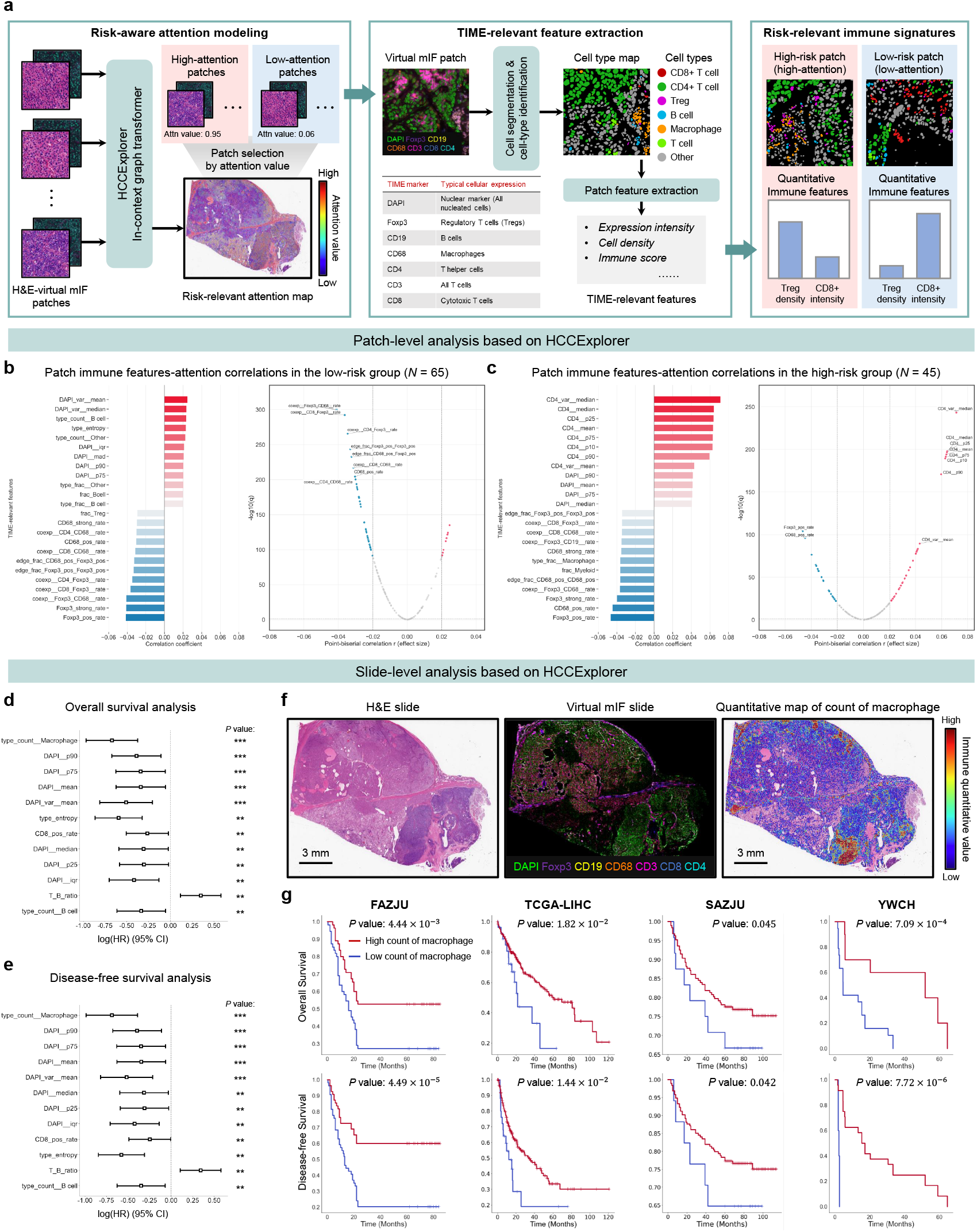
HCCExplorer captures biologically interpretable immune features and identifies macrophage infiltration as a robust prognostic determinant. **a** Schematic illustration of the interpretability framework. The workflow links the model’s risk-aware attention scores from the graph transformer to quantifiable TIME-relevant features. By performing cell segmentation on high-attention vs. low-attention patches, the model translates abstract computational weights into tangible biological signatures. Note that CD4+ T cells represent Foxp3-negative helper T cells, while Tregs represent CD4+Foxp3+ cells. **b, c** Identification of risk-specific immune signatures. Bar plots (left) and volcano plots (right) display the correlation between patch-level attention scores and quantitative immune features in the **b** low-risk group and **c** high-risk group. Red bars indicate a positive correlation (features prioritized by the model), while blue bars indicate a negative correlation. **d, e** Screening for prognostic biomarkers. Forest plots summarizing univariate Cox proportional hazards regression analysis for **d** OS and **e** DFS. Features are ranked by Log hazard ratio (HR). Macrophage density per patch (type_count_of_macrophage) emerges as the most significant protective factor, while T/B cell ratio is a significant risk factor. Wald test was used to calculate *P* values for each covariate. **f** Spatial visualization of the top-ranked biomarker. Representative images showing an H&E whole-slide (left), its corresponding generated virtual mIF slide (middle), and a quantitative heatmap of macrophage count (right). **g** Cross-cohort validation of macrophage stratification. Kaplan-Meier survival curves for OS (top) and DFS (bottom) stratified by HCCExplorer-derived macrophage count. The stratification capability is validated across four independent cohorts: FAZJU (internal), TCGA-LIHC (external), SAZJU (external), and YWCH (external). High macrophage count per patch (red line) is consistently associated with significantly better prognosis across all centers (Log-rank test *P* values are indicated). Significance levels for this figure: * *P* < 0.05, ** *P* < 0.01, *** *P* < 0.001.

We first investigated which immune features dominated the model’s decision-making process by analyzing the correlation between patch attention scores and quantitative immune features (Figure 4b, c). In the low-risk group, the model prioritized features indicative of active anti-tumor immunity, showing significant positive correlations with B-cell density (type_count_B cell) and immune entropy (type_entropy), indicating that diverse immune infiltration is a key driver for a favorable prognosis (Figure 4b). Conversely, in the high-risk group, the model’s focus shifted toward T-cell dysregulation, with the variance of CD4 signal intensity (CD4_var_median) showing the strongest association, indicating that heterogeneous CD4 expression is a hallmark of aggressive disease (Figure 4c, Supplementary Information Table S3).

To validate the clinical relevance of these attention-derived features, we screened the top biomarkers using univariate Cox regression for OS and DFS (Figure 4d, e). Most notably, Macrophage density per patch (type_count_Macrophage) emerged as the dominant prognostic factor, exhibiting the strongest protective effect in both OS (Hazard Ratio = 0.51, 95% CI: 0.38– 0.69, *P* < 0.001) and DFS (Hazard Ratio = 0.50, 95% CI: 0.37–0.68, *P* < 0.001) (Figure 4d, e, Extended Data Figure 6). The predictive potency of this feature surpassed that of well-established immuno-oncology paradigms also validated in our cohort, such as CD8+ T cell positivity^43^ (OS Hazard Ratio = 0.77; DFS Hazard Ratio = 0.78) and the T/B cell ratio^44^ (OS Hazard Ratio = 1.42; DFS Hazard Ratio = 1.41). Given that macrophage count was identified as the top-ranked protective feature, we further validated its prognostic utility at the slide level (Figure 4f, Extended Data Figure 6a, Methods “Immune feature analysis”). We visualized the spatial distribution of this biomarker, generating a quantitative map of macrophage counts that highlighted the pronounced intratumoral heterogeneity captured by HCCExplorer, in which regions of dense macrophage infiltration corresponded to the model’s high-attention, low-risk predictions (Figure 4f, Extended Data Figure 6a).

Finally, to assess the clinical generalizability of this model-discovered biomarker, we performed Kaplan-Meier analysis across four cohorts (Figure 4g). Stratification based on the HCCExplorer-derived macrophage count yielded consistent and significant separation of patient outcomes. High macrophage count was significantly associated with better OS (*P*=4.44 × 10^−3^) and DFS (*P* = 4.49 × 10 ^−5^) in the internal FAZJU cohort. Crucially, this prognostic power was robustly reproduced in external datasets, including TCGA-LIHC (OS: *P =* 0.018), SAZJU (OS: *P* =0.045), and notably the YWCH cohort, where stratification was exceptionally distinct (OS: *P =* 7.09 × 10^−4^ ; DFS: *P=* 7.72 × 10^−6^) (Figure 4g). To further evaluate the predictive stability of this biomarker, we compared its C-index with that of the full HCCExplorer model (Table 2, Extended Data Table 1, Methods “Immune feature analyses”). As expected, the single-feature macrophage abundance yielded a slightly lower C-index in the internal FAZJU cohort (0.65, 95% CI: 0.56–0.75) compared to the comprehensive multi-feature HCCExplorer (0.71, 95% CI: 0.66–0.74), reflecting the latter’s capacity to capture complex local patterns. However, in the external YWCH cohort, the macrophage biomarker outperformed the full model (C-index: type_count_Macrophage 0.62 (95% CI: 0.52–0.72) vs. HCCExplorer 0.52 (95% CI: 0.50–0.55)). This inversion underscores that, although full graph neural networks capture comprehensive spatial features, the specific macrophage infiltration density identified by HCCExplorer constitutes an effective biological signal with robust generalization across heterogeneous datasets. These results demonstrate that HCCExplorer successfully identifies and prioritizes biologically distinct features, specifically macrophage infiltration density, establishing it as a robust, cross-cohort prognostic biomarker directly from standard H&E slides (Figure 4, Extended Data Figure 6b, Extended Data Table 1).

### 5 Multi-modal integration reveals the protective role of M1-polarized macrophages in hepatocellular carcinoma

Having established macrophage infiltration as a robust prognostic determinant (Figure 4), we sought to dissect the molecular underpinnings of this protective effect. To achieve this, we developed a multi-modal integration strategy that bridges morphological patterns with molecular signatures (Figure 5a). We combined HCCExplorer, which captures the spatial distribution and density of macrophages within the tumor microenvironment, with CIBERSORTx^45^ to deconvolute bulk RNA sequencing (RNA-seq) data and estimate the relative abundances of fine-grained immune cell subtypes on the TCGA-LIHC cohort (Figure 5b, c, Methods “Cell-type deconvolutions of bulk RNA-seq”).

**Figure 5.**
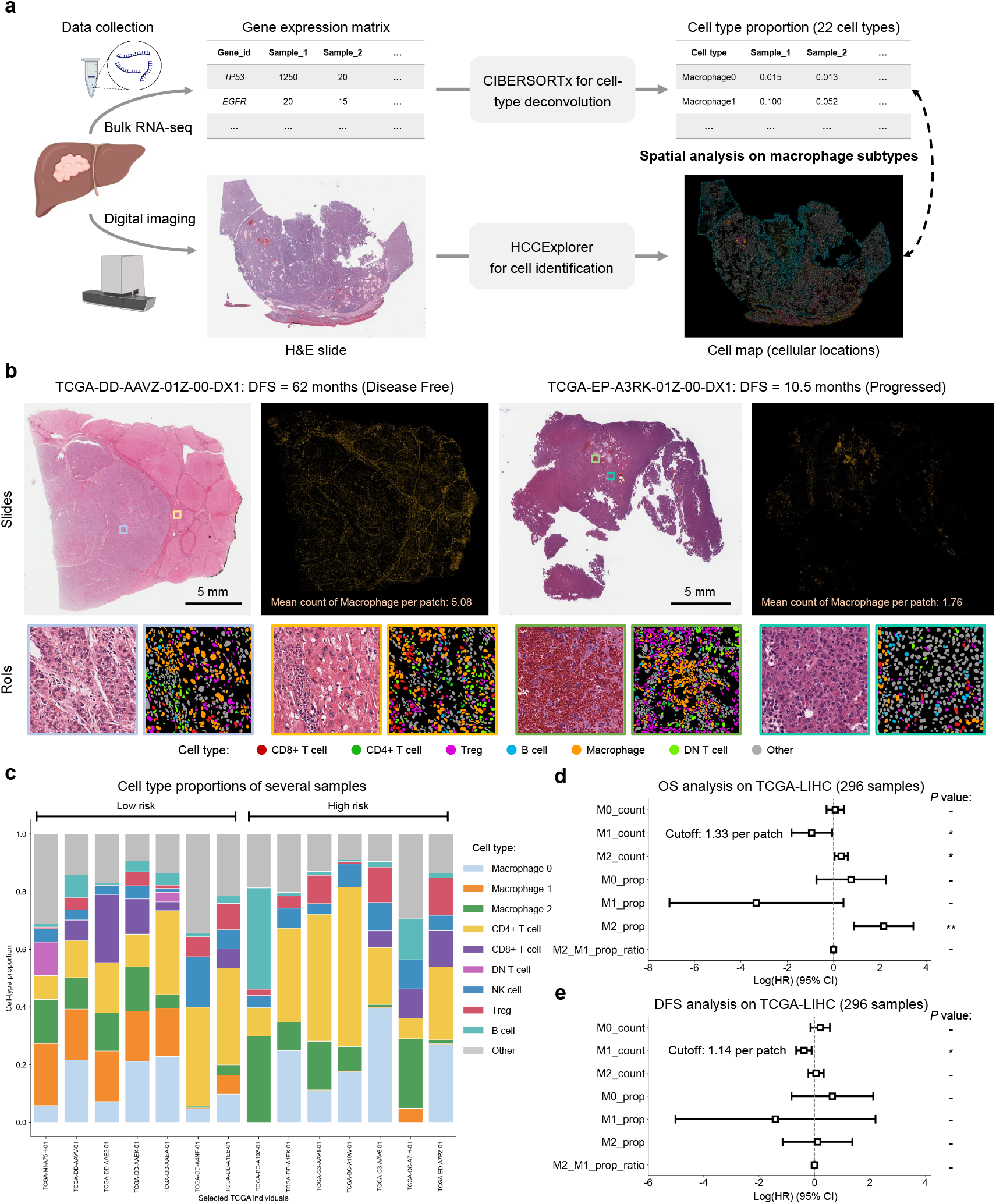
Multi-modal integration reveals the protective role of M1-polarized macrophages in HCC. **a** Schematic of the multi-modal integration strategy. The workflow illustrates the methodology for bridging spatial pathology and transcriptomics. It combines the spatial cellular density derived from HCCExplorer (absolute cell counts per patch) with immune cell proportions inferred from bulk RNA-sequencing to estimate the absolute density of specific macrophage subtypes (e.g., M1 vs. M2). **b** Representative visualization of immune phenotypes. Comparison between a long-term survivor (left) and a rapid progressor (right) from the TCGA-LIHC cohort. The panel displays the whole-slide H&E image, the generated virtual mIF map focusing on macrophage distribution (yellow signals), and magnified RoIs showing the cell classification results. The survivor exhibits a “hot” phenotype with dense, diffuse macrophage infiltration (Mean count: 5.08), whereas the progressor shows a “cold” phenotype with sparse infiltration (Mean count: 1.76). **c** Immune composition profiling. Stacked bar plots displaying the relative proportions of deconvoluted immune cell subsets (including Macrophage M0, M1, M2 subtypes) across representative low-risk and high-risk individuals. **d, e** Prognostic assessment of macrophage polarization. Forest plots summarizing univariate Cox regression analysis for **d** overall survival and **e** disease-free survival in the TCGA-LIHC cohort (296 patients used here as they have both HCCExplorer and cell-type deconvolution results). The plots evaluate the hazard ratios for both the derived absolute counts (“_count”) and relative proportions (“_prop”) of macrophage subtypes. The analysis reveals a functional dichotomy: the absolute density of M1 macrophages (M1_count) is a significant protective factor, whereas M2 prevalence (M2_prop) and the M2/M1 ratio are risk factors for poor prognosis. Error bars represent 95% CI. Wald test *P* values are indicated, significance levels: * *P* < 0.05, *P* < 0.01**, *** *P* < 0.001.

By synthesizing these inputs to calculate the absolute density of specific macrophage polarization states^46^, the analysis revealed a critical functional dichotomy (Figure 5d, e). The absolute density of M1 macrophages (count of M1 macrophages per patch), characterized by their pro-inflammatory and tumoricidal activity^47^, emerged as a significant protective factor for both OS and DFS (OS: Hazard Ratio = 0.40, 95% CI: 0.16–0.94, *P* < 0.05; DFS: Hazard Ratio = 0.68, 95% CI: 0.52–0.90, *P* < 0.05). In contrast, markers associated with M2 macrophages, such as the proportion of M2 cells, were identified as risk factors^48^ associated with poor overall survival (Hazard Ratio = 8.72, 95% CI: 2.42–31.39, *P* < 0.01). Collectively, these findings uncover an immune feature in HCC, demonstrating that the favorable prognosis associated with macrophage infiltration is not generic but is predominantly driven by the dense accumulation of anti-tumor M1-like subtypes. This specific biological signal, successfully identified by HCCExplorer with CIBERSORTx from standard H&E slides and bulk RNA-seq, underscores the critical prognostic value of resolving macrophage heterogeneity.

### 6 HCCExplorer identifies macrophage enrichment at the invasion frontier as a critical protective barrier

While patch-level analysis identified macrophage density as a prognostic factor, it does not fully account for the complex spatial organization of the TIME^49^. To decipher the spatial architecture of immune interactions, we leveraged the graph transformer^50^ module embedded within HCCExplorer (Figure 6a). This design enables the model to capture long-range cellular cross-talk and structural motifs beyond the field of view of a single patch (Figure 6a, Extended Data Figures 2a, 7, and 8, Methods “Graph-based multi-modal contextual survival analyses” and “Immune feature analyses”).

**Figure 6.**
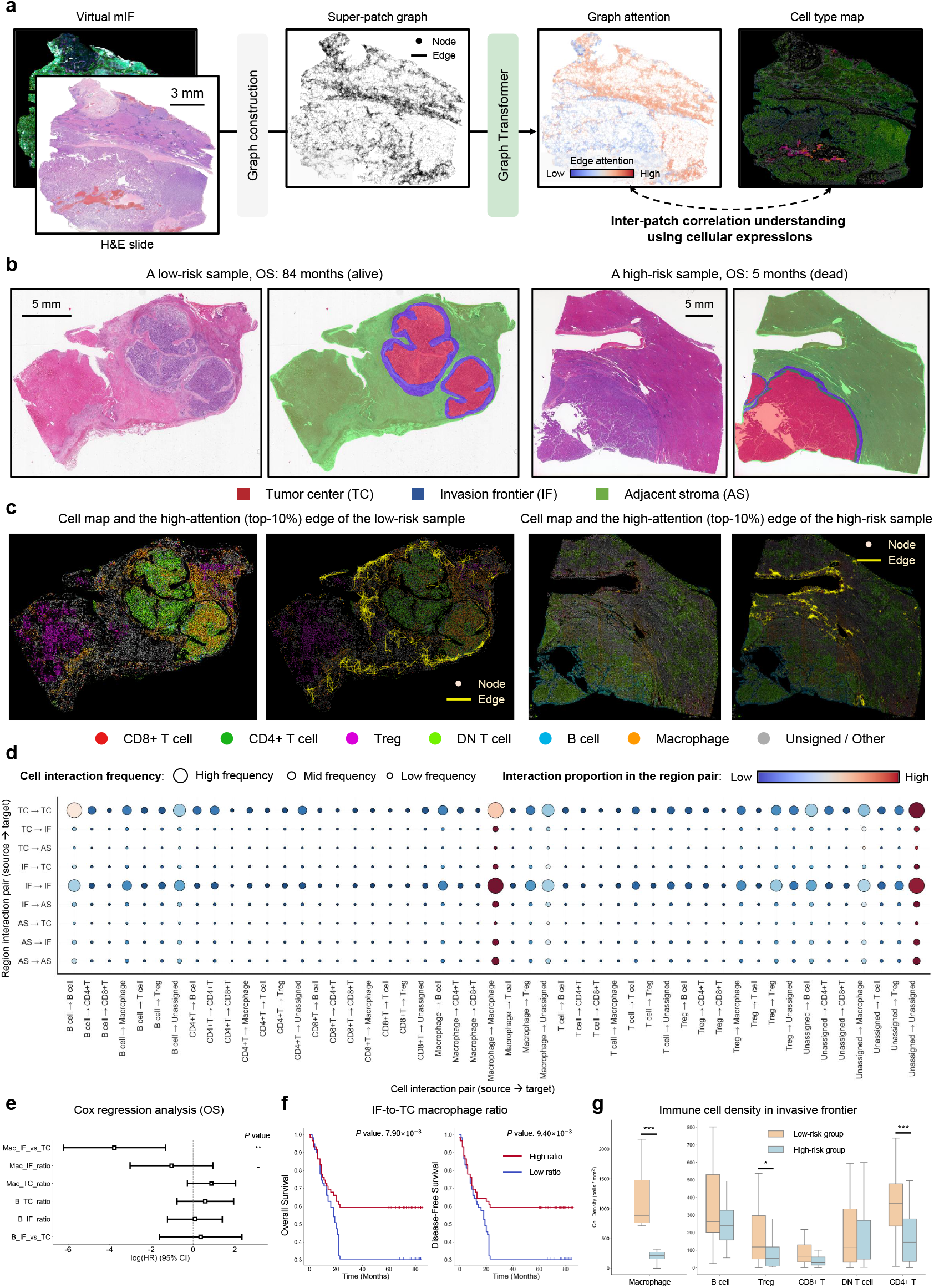
Graph-based spatial modeling in HCCExplorer reveals macrophage dynamics at the invasion frontier. **a** Workflow of the inter-patch correlation understanding using HCCExplorer. Virtual mIF slides are converted into a graph structure where nodes represent tissue super-patches. The model learns global context via attention mechanisms to generate a cell type map that incorporates inter-patch correlations. **b** Spatial segmentation of WSIs into tumor center (TC, red), invasion frontier (IF, blue), and adjacent stroma (AS, green), shown for representative low-risk and high-risk patients. **c** Visualization of the graph transformer. The overlay shows the high-attention edges (top 10% weights), revealing dense communication networks anchored at the invasion frontier in the low-risk sample. **d** Bubble plot of region-specific cellular interactions. Rows represent directional interactions between spatial regions (e.g., IF to TC), and columns represent cell-cell pairs. Circle size indicates frequency; color intensity indicates the proportion of interaction. Interactions involving macrophages originating from the invasion frontier are dominant. Notably, in this analysis, graph nodes represent high-attention image patches, and “cell type” refers to the dominant cell phenotype within each patch, serving as a proxy for the local cellular microenvironment. **e** Forest plot of Cox regression for spatial metrics. The Macrophage IF-to-TC ratio (Mac_IF_vs_TC) is identified as a significant protective factor. Wald test *P* values were reported. **f** Kaplan-Meier survival curves for OS and DFS stratified by the IF-to-TC macrophage ratio. High ratio (red) correlates with better outcomes. Log-rank test was conducted. **g** Comparative analysis of the immune landscape at the invasion frontier. Box plots showing the densities of major immune subsets in the low-risk group (10 representative patients) versus the high-risk group (10 representative patients). Mann-Whitney U Test was used for statistical analysis. Significance levels: * *P* < 0.05, ** *P* < 0.01, *** *P* < 0.001.

To interpret these learned spatial dependencies, we stratified WSIs into three biologically distinct compartments: the tumor center (TC), the invasion frontier (IF), and the adjacent stroma (AS) (Figure 6b). We subsequently mapped the graph attention weights onto these regions to identify the spatial interactions prioritized by the model for risk prediction (Figure 6c). Visualization of the “high-attention” edges revealed a striking pattern: in low-risk patients, the model consistently highlighted dense connectivity networks anchored at the invasion frontier, suggesting that immune activity at the tumor boundary is more determinative of clinical outcomes than that within the tumor core (Figure 6b, c).

We then quantified this by calculating the interaction frequency between different regions and cell types (Figure 6d). The bubble plot of region-cell interactions revealed that the invasion frontier (IF) acts as the central hub of immune communication. Specifically, pathways originating from the IF (e.g., IF-to-IF) exhibited the highest interaction proportions. Crucially, dissecting the cellular composition of these interactions showed that Macrophages were the dominant drivers. The macrophage-to-macrophage within the IF and from IF to TC represented the most frequent and specific communication motifs, far surpassing B- or T-cell interactions in this spatial context.

Based on the explicit focus of the model on IF-driven macrophage interactions, we hypothesized that the spatial distribution of macrophages, specifically their accumulation at the IF relative to the TC, serves as a critical biomarker. To quantify this spatial heterogeneity, we defined a metric: the IF-to-TC Macrophage Ratio. We screened multiple spatial metrics using Cox regression analysis, and the IF-to-TC Macrophage Ratio emerged as the most significant protective factor (OS: Hazard Ratio = 0.02, 95% CI: 0.00–0.26, *P* < 0.01), outperforming B-cell-related spatial metrics (Figure 6e). This stratification was further corroborated by Kaplan-Meier analysis, which demonstrated that patients with a high IF-to-TC macrophage ratio exhibited significantly superior OS (*P* = 7.90 × 10^−3^) and DFS (*P*= 9.40 × 10^−3^) compared to those with a low ratio (Figure 6f).

To elucidate the mechanistic basis of this “Macrophage Barrier,” we examined the broader cellular and morphological landscape at the invasion front (Figure 6g, Extended Data Figure 9). The analysis revealed that this phenomenon extends beyond simple macrophage accumulation and represents a coordinated “Containment Niche” in low-risk cases. Upon histopathological review, low-risk cases with high macrophage infiltration frequently exhibited a desmoplastic, encapsulated growth pattern, surrounded by a fibrotic rim (Extended Data Figure 9a), in contrast to the infiltrative or budding patterns observed in high-risk, macrophage-depleted cases (Extended Data Figure 9b). These observations suggest that the enriched macrophages likely contribute to active matrix remodeling, recruiting myofibroblasts to form a fibrous capsule that physically constrains tumor expansion. Notably, within this niche, we observed a distinct co-localization triad of CD68+ macrophages, CD4+ helper T cells, and Foxp3+ Tregs (Figure 6g, Extended Data Figure 9a). While Tregs are typically associated with immunosuppression and poor outcomes, their specific enrichment at this desmoplastic rim implies a context-dependent protective role, potentially by maintaining the stability of this architectural barrier (Extended Data Figure 9a), a phenomenon analogous to favorable Treg infiltration at the invasion front in colorectal cancer^51,52^. Collectively, these findings suggest that HCCExplorer has identified a critical “Containment Niche”, where the strategic co-localization of macrophages and T cells at the tumor edge orchestrates a desmoplastic response to effectively wall off the tumor, thereby preventing invasion and conferring a survival advantage (Figure 6, Extended Data Figure 9).

## Discussion

The fundamental limitation of routine hepatopathology is its inability to resolve the complex tumor immune microenvironment (TIME), thereby rendering critical prognostic information inaccessible. Although recent advances like GigaTIME^25^ and HEX^26^ have demonstrated promise in H&E-to-protein prediction, maintaining cellular fidelity remains a significant challenge in the absence of pixel-aligned supervision. HCCExplorer addresses this issue by introducing the Cell Consistent Cross-Modal Unpaired Translation (C3UT) model. By enforcing cellular topology constraints derived from histology images, the model suppresses hallucination artifacts frequently observed in generative adversarial networks. This ensures that synthesized biomarkers are not only statistically plausible but also structurally accurate at the single-cell level. The robustness of this biologically constrained approach is highlighted by its data efficiency. Despite training on a curated dataset of only 30 paired slides, HCCExplorer successfully generalized to a large-scale multi-center cohort of 1,813 patients comprising approximately 9 billion cells. This high-fidelity reconstruction provides a necessary biological basis for downstream analysis and prevents the learning of spurious correlations. Consequently, HCCExplorer outperforms both pathology and proteomics foundation models in prognostic stratification, underscoring the importance of explicit biological modeling for medical AI.

Beyond technical innovation, the proposed framework resolves existing controversies regarding the role of macrophages in hepatocellular carcinoma (HCC). Although tumor-associated macrophages are frequently categorized as immunosuppressive drivers of progression, the spatially resolved analysis in this study identifies macrophage infiltration as a primary protective determinant. Through multi-modal integration with bulk RNA-seq, we disentangled this signal, revealing that the survival benefit is predominantly driven by M1-polarized phenotypes rather than the pro-tumorigenic M2 subset. This finding highlights the capacity of virtual immunophenotyping to capture functional dichotomies that are typically obscured in bulk tissue analysis.

A significant biological insight enabled by graph-based multi-modal contextual learning is the identification of a coordinated “Containment Niche” at the invasion frontier of hepatocellular carcinoma. We observed that low-risk patients exhibit a dense accumulation of macrophages and T cells, which form a desmoplastic rim that physically constrains tumor expansion. Notably, the model revealed that Foxp3+ Tregs are enriched within this specific spatial compartment. In contrast to the generally immunosuppressive role observed in the tumor core, the co-localization of Tregs with macrophages at the invasion frontier suggests a context-dependent protective function. We hypothesize that these Tregs may contribute to stabilizing the architectural integrity of the containment barrier or to dampening pro-invasive inflammatory responses in the peritumoral stroma. This observation aligns with emerging evidence in other solid tumors^53,54^ suggesting that the prognostic impact of immune cells is fundamentally determined by their spatial neighborhood.

Several limitations of this study warrant consideration. First, while virtual staining provides spatial resolution for major cell lineages, discerning subtle functional states, such as macrophage polarization, currently requires integration with bulk transcriptomics. Future iterations will aim to predict functional markers directly to achieve higher granularity. Second, although validated across multiple independent cohorts, the retrospective nature of the study necessitates prospective evaluation to confirm clinical utility. Moving forward, integrating HCCExplorer into clinical trials could provide real-time stratification to guide adjuvant therapies.

In conclusion, HCCExplorer establishes a practical pathway for next-generation computational pathology in hepatocellular carcinoma. By extracting interpretable, spatially resolved immunophenotypes from routine slides, we uncovered the macrophage-mediated containment barrier and the distinct spatial role of Tregs at the invasion frontier. While the primary validation focuses on liver cancer, the underlying computational workflow is inherently generalizable. We anticipate that this pipeline will be readily adaptable to other solid tumors, offering a scalable tool to decode universal spatial immune rules across the pan-cancer landscape. Ultimately, this work validates the paradigm that artificial intelligence can transform standard medical imaging into a rich source of biological insight.

## Methods

### 1 Cohorts used in HCCExplorer

To develop and validate HCCExplorer, we curated a large-scale, multi-center retrospective study comprising a total of 1,813 patients with hepatocellular carcinoma (HCC) from four independent institutions, namely, The First Affiliated Hospital, Zhejiang University School of Medicine (FAZJU), The Second Affiliated Hospital, Zhejiang University School of Medicine (SAZJU), The Cancer Genome Atlas - Liver Hepatocellular Carcinoma^55^ (TCGA-LIHC), and Yiwu Central Hospital (YWCH). Clinicopathological characteristics of the study population are summarized in Table 1. To train HCCExplorer, the primary cohort from FAZJU was randomly partitioned into a training set (949 patients, 1,017 H&E slides, 30 mIF slides) for model development and an internal test set (237 patients, 237 H&E slides, 10 mIF slides) for initial evaluation. To assess the model’s generalizability across diverse populations, we utilized three independent external validation cohorts: SAZJU (211 patients, 211 H&E slides), YWCH (74 patients, 74 H&E slides), and the publicly available TCGA-LIHC dataset (342 patients, 342 slides). The study cohorts covered a broad spectrum of clinical profiles, ensuring representative coverage of patient heterogeneity. Patient ages ranged from 28 to 89 years, with a predominance of male patients consistent with HCC epidemiology^56^ (Table 1). The cohorts encompassed comprehensive histological stratifications, including tumor multiplicity (single vs. multiple), AJCC stage (I–III), and differentiation grade (I–IV). Furthermore, robust follow-up data were available across cohorts, with documented overall survival (OS), disease-free survival (DFS), and recurrence-free survival (RFS) times, providing a solid foundation for survival analysis.

### 2 H&E-to-virtual mIF translation

To enable the generation of spatially resolved proteomic data from standard histology, we developed the Cell-Consistent Cross-Modal Unpaired Translation (C3UT) model to synthesize virtual multiplex immunofluorescence (mIF) images directly from H&E-stained images (Figure 1a).

#### 2.1 Dataset construction and quality control

We collected 40 pairs of H&E and multiplex immunofluorescence (mIF) whole-slide images (WSIs) from the internal FAZJU cohort, with 30 pairs for training and 10 pairs for independent testing. Each mIF image contains 8 channels, DAPI, CD3, CD4, CD8, CD19, CD68, Foxp3, and Sample Auto-Fluorescence (SampleAF). Notably, the H&E and mIF images were derived from consecutive serial tissue sections. While we used the pathology registration framework VALIS^57^ for fine slide-level registration, pixel-to-pixel alignment remained unachievable due to intrinsic morphological deformations and tissue variability across serial sections (Extended Data Figure 1a). Consequently, we adopted an unpaired image translation strategy for the Cell-Consistent Cross-modal Unpaired Translation (C3UT) model to accommodate these spatial discrepancies. To ensure robust model optimization, we implemented a rigorous quality control (QC) pipeline to maximize the informational density specifically for the training set. Initially, 219,416 coarsely aligned patches of 1024 × 1024 pixels were extracted from the 30 training slide pairs at 20 × magnification. We then applied a dual-modality filtering mechanism based on cell segmentation results, in which HoVerNet^58^ was used for H&E cell segmentation, and StarDist^59^ for mIF. A patch pair was only retained if meaningful biological signals were detected in both the H&E (sufficient cellularity) and mIF (sufficient marker expression) domains. This strict filtering effectively eliminated low-quality regions—such as background, necrosis, or tissue folding— yielding a high-quality training dataset of 154,916 patch pairs covering markers including CD3, CD4, CD8, CD19, Foxp3, CD68, DAPI, and corresponding SampleAF. For comprehensive evaluation, the 10 withheld independent slide pairs, comprising a total of 43,618 H&E-mIF patch pairs, were employed to assess the H&E-to-mIF generation performance through both quantitative analysis and a visual Turing test (specific assessment protocols are detailed in the Methods “Evaluation metrics” section).

#### 2.2 Cell-consistent cross-modal unpaired translation model

To translate H&E images (*X*) into corresponding virtual multiplex immunofluorescence (mIF) images (*Y*) without requiring pixel-perfect paired data, we developed the C3UT model. Our architecture is a generative adversarial network^60^ (GAN) built on the Contrastive Unpaired Translation^34^ (CUT) framework, but it introduces specific architectural and input-level constraints to preserve biological fidelity (Extended Data Figure 1b).

##### 2.2.1 Network architecture

The generator (*G*) employs a ResNet-based architecture with 9 residual blocks^61^ (each block contains 2 layers), designed to preserve high-frequency morphological details during the domain transformation. For the discriminator (*D*), we utilized a 3-layer PatchGAN^62^, which classifies local 70 × 70-pixel patches as real or fake, emphasizing high-frequency texture consistency.

##### 2.2.2 Channel-wise input-output formulation

Standard stain-to-stain translation often maps 3-channel H&E to a single-channel marker density. However, to better model the intrinsic spatial dependency between protein expression and cellular localization, we designed a multi-channel target domain. For each specific marker translation task, the model maps the 3-channel H&E input to a 3-channel composite output: [DAPI, Marker, SampleAF]. By explicitly predicting the DAPI (cellular signal) and SampleAF channels alongside the target marker, the model is forced to adaptively perceive the relationship between cellular positioning and marker expression, thereby reducing false-positive signals in acellular regions.

##### 2.2.3 Cell-map-guided adversarial and contrastive learning

To address semantic hallucination that is common in unpaired translation, C3UT introduces a cell-map-guided adversarial constraint. The discriminator is conditioned on a binary cell map *M*_*x*_, derived from the H&E input via a cell segmentation network, HoVerNet^58^ (a state-of-the-art model optimized for separating overlapping nuclei in H&E slides), assessing the joint probability of the generated image and its underlying cellular distribution, where “1” means cellular region and “0” is background. Conversely, for the real mIF image, the cell map *M*_*y*_ is extracted by StarDist from the DAPI channel, leveraging its superior performance in detecting star-convex nuclear shapes within fluorescence intensity fields. This domain-specific selection ensures high-fidelity cellular localization for both modalities. The above process can be described as:

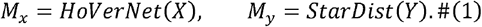

This ensures that synthesized signals physically align with existing cells. Simultaneously, to maintain semantic correspondence, we employed a multilayer contrastive loss^34^ (PatchNCE). Features were extracted from five specific layers of the generator encoder (indices 0, 4, 8, 12, and 16) to capture a hierarchy of representations ranging from low-level textures to high-level semantics. This loss ℒ_contrastive_ maximizes the mutual information between corresponding patches in the H&E input *X* and virtual mIF output Ŷ= *G (X)* The multilayer contrastive loss, based on the PatchNCE^34^ framework, can be expressed as:

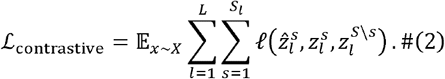

The contrastive loss ℓ (·) is defined as:

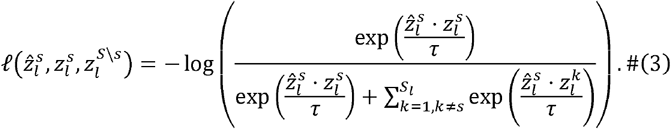

Here, *l* ∈ {0,4,8,12,16} denotes the indices of selected layers within the encoder. *s* ∈ {1,…,*S*_*l*_ } is the spatial index representing a specific patch location within the feature map of layer *l*. 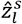 (Query) is the feature vector at spatial location in layer *l* of the synthesized image *G*(*X*), serving as the anchor or query sample.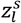 (Positive) is the corresponding feature vector at the same spatial location *s* in layer *l* of the input image *x*. This serves as the positive sample, enforcing semantic consistency between the input and output. 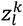(Negatives) are the feature vectors at other spatial locations *k* (where *k* ≠ *s*) within the input image *x*. These serve as negative samples to be pushed away from the query. τ is the temperature hyper-parameter, empirically set to 0.07, which scales the logits to control the smoothness of the distribution.

The total objective of C3UT is formulated as a weighted sum of the multilayer contrastive loss and the cell-guided adversarial loss. To enhance training stability and mitigate the vanishing-gradient problem, we adopt the Least Squares GAN^63^ (LSGAN) framework rather than the standard negative log-likelihood. The adversarial component is designed to ensure that the synthesized mIF images follow the distribution *D* of the target domain while maintaining spatial consistency guided by cell maps. The discriminator is trained to distinguish between real image-mask pairs and synthesized pairs by minimizing the following objective:

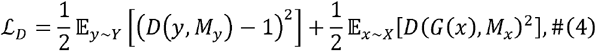

where *M*_y_ and *M*_x_ represent the cell maps derived from the real target image *y* and the source H&E image *x*, respectively. The generator *G* aims to produce realistic mIF images *G* (*x*)to confuse the discriminator. The adversarial loss for the generator is defined as:

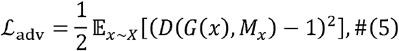

The final objective function for C3UT is:

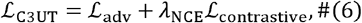

where *λ*_NCE_ controls the relative importance of the contrastive term. In our experiments, we set λ_NCE_ =1 in the HCCExplorer training protocol.

##### 2.2.4 Training configuration

The training images were initially loaded at a resolution of 1024 × 1024 pixels (20 × magnification) and subsequently resized to 512 × 512 pixels to optimize computational efficiency while preserving sufficient cellular resolution. Following the settings of CUT^34^, we set the batch size to 1 and an Adam optimizer with a learning rate of 0.0002. For each marker, we trained 10 epochs (around 1,550,000 steps) on an NVIDIA GeForce RTX 4090 24G GPU and used the final weights for testing.

### 3 H&E-virtual mIF contrastive learning

To effectively fuse the morphological information from H&E images with the molecular insights from the virtual mIF data, we developed a multi-modal contrastive learning framework based on a multi-stain pretraining algorithm MADELEINE^64^ (Figure 1b, Extended Data Figure 2a-c). The co-optimization framework aligns the representations of H&E slides with their corresponding virtual mIF counterparts at both global (slide-level) and local (patch-level) scales.

#### 3.1 Multi-modal feature encoding with marker-specific adapters

First, paired real H&E and generated virtual mIF WSIs were tiled into non-overlapping patches of 256 × 256 pixels at a resolution of 0.5 *μm*/pixel (Extended Data Figure 2a) using TRIDENT^65^. To extract high-level feature representations, we employed a pre-trained Vision Transformer^66^ (ViT), UNI-h2^29^, as the backbone encoder. The backbone weights were frozen to maintain robust generalizability.

Distinct from previous approaches that utilize simple stain embeddings, we introduced trainable marker-specific adapters to capture the unique biological semantics of individual channels. For each input modality/marker (H&E, DAPI, CD3, CD4, CD8, CD19, CD68, and Foxp3), the frozen ViT features are passed through a dedicated adapter network—a lightweight multilayer perceptron (MLP) including two fully connected (FC) layers with a ReLU^67^ activation layer— which projects the generic visual features into a marker-specific latent space (from 1,536 to 256 dimensions). Specifically, let 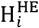 denote the adapted patch features for the H&E modality (and analogously for other stains). Subsequently, these patch embeddings are aggregated into slide-level representations using a multi-head gated attention network^40^ with *M* heads. The attention weights for the *m*-th head, denoted as 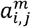, are defined as:

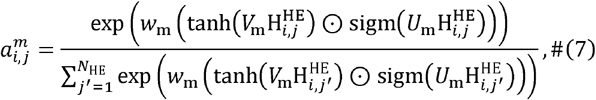

where 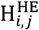 is the *j*-th patch feature for the *i*-th slide.

Finally, the outputs of all attention heads are concatenated and projected via a linear layer to yield the final slide embedding 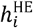.

#### 3.2 Global cross-modal alignment

To learn a unified patient representation, we applied a global objective to align slide embeddings from the H&E modality with each corresponding virtual mIF marker in a shared latent space (Extended Data Figure 2b). Following the MADELEINE^64^ framework, we employed a symmetric cross-modal contrastive learning objective^68^ ℒ_InfoNCE_ . For a batch of *B* cases, each containing *K* stain pairs, the loss is defined as:

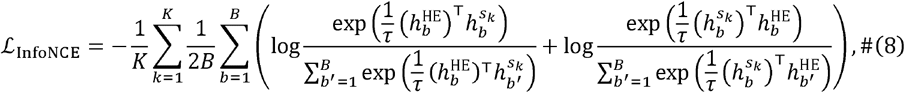

where *s*_*k*_ represents a non-H&E stain, τ represents the Softmax temperature hyper-parameter, which is set as 0.07 in the work. This objective enforces that slide embeddings from the same patient case (positive pairs) are closer to each other while pushing embeddings from unpaired slides (negative pairs) apart within the batch. Each term maximizes the dot-product similarity between embeddings from the same pair, normalized by negative similarity pairs, ensuring that the model captures globally consistent phenotypic patterns across modalities.

#### 3.3 Local graph optimal transport alignment

To enforce fine-grained correspondence and avoid reliance solely on global statistics, we incorporated a local alignment mechanism based on Graph Optimal Transport^69,70^ (GOT) (Extended Data Figure 2c). Formally, we define the empirical distribution 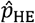 of the H&E patch embeddings as:

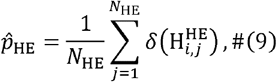

where *N*_HE_ is the number of H&E patches, δ(·) denotes the Dirac-delta function, 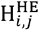 is the embedding of the *j*-th H&E patch. Correspondingly, the *k*-th marker’s empirical distribution 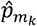 can be formulated as:

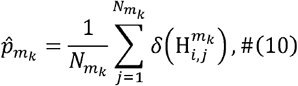

where 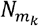 is the number of the *k*-th marker’s patches and 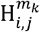 is the embedding of the *k*-th marker patch.

We construct an H&E graph: 𝒢_HE_ (*V*_HE_, *E*_HE_), where edges are formed based on cosine similarity thresholds. We align the stain-specific graphs by minimizing two metrics:

(i) Wasserstein Distance (ℒ_Node_): Defined between the empirical distributions of patch embeddings (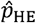 and 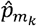). Intuitively, this computes the distance between the node embedding distributions, ensuring that local morphological features are correctly mapped:

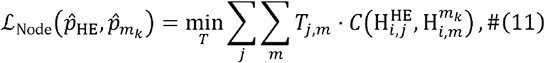

where *T* denotes the transport plan, *m*_k_ is the *k*-th marker, and *C*(·, ·) represents the cost function between embeddings using the cosine distance metric.

(ii) Gromov-Wasserstein Distance (ℒ_Edge_): Defined between the edges of 𝒢_HE_ and 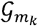. This enforces the graphs to follow a similar structure (or topology), preserving the spatial organization of the tissue microenvironment:

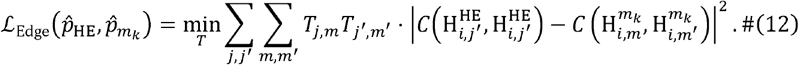

The local alignment objective ℒ_GOT_ is given as the combination of these two metrics over all *K* markers, with *γ* denoting a weighting term (γ is set to 0.5 in HCCExplorer):

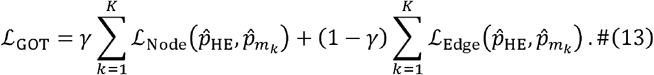

The final training objective ℒ_co_ combines the global and local losses:

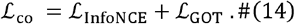

We performed this co-optimization training using the internal training set from the FAZJU cohort, comprising 949 patients and 1,027 paired H&E and virtual mIF slides. The model was trained for 200 epochs with the batch size of 64 on 8 NVIDIA GeForce RTX 4090 24G GPUs. The detailed settings are identical to those of MADELEINE^64^. Upon completion, the optimized H&E adapter and the specific mIF channel adapters were preserved and frozen. These fine-tuned adapters serve as specialized feature extractors to generate node embeddings for the graph neural network (GNN) modeling and survival analysis.

### 4 Graph-based multi-modal contextual survival analyses

To decode the complex spatial interactions within the TIME for prognostic prediction, we developed a graph-based deep learning framework that integrates the learned H&E and virtual mIF representations (Figure 1c, Extended Data Figure 2d).

#### 4.1 Graph construction for H&E-virtual mIF slides

We first segment the H&E slide into non-overlapping patches of 256 × 256 pixels at 20 × magnification. Patches containing more than 75% background are discarded. Each remaining patch is encoded into a 256-dimensional feature vector using the pre-trained UNI-h2^29^ encoder with H&E-specific feature adapter. To reduce the computational burden of processing a vast number of patches while preserving local histological homogeneity, we aggregate patches into “super-patches” following TEA-Graph^71^. Specifically, spatially adjacent and morphologically similar patches are merged based on two simultaneous criteria: (i) the cosine similarity of their patch-level features, and (ii) the Euclidean distance of their spatial coordinates. Patches satisfying both criteria are grouped into the same super-patch. In cases where a patch meets the merging criteria for multiple potential groups, it is assigned to the super-patch containing the largest number of constituent patches. Subsequently, the feature representation of each super-patch is computed as the mean of the features of all its member patches. Graph edges are then defined based on the spatial distance between these super-patches.

Building on the H&E-based graph, we extend this framework to mIF slides. Using the virtual mIF slide with the H&E slide, we directly reuse the spatial coordinates and super-patch boundaries defined in the WSI graph. This design ensures spatial consistency across modalities, enabling precise alignment between H&E super-patches and their corresponding IF counterparts for joint graph-based analysis.

#### 4.2 Polynomial node feature optimization in multi-modal graphs

To enhance contextual learning, we adopt a graph-based approach tailored for modeling both the structural topology and the constituent patches within a WSI. We construct a multi-modal graph to integrate information from H&E and mIF domains. Specifically, H&E nodes are derived from the spatially defined super-patches, while mIF nodes correspond to the same super-patch regions across multiple immune channels. To capture cross-modal dependencies, edges are established between neighboring H&E nodes to capture local tissue context, and between each H&E node and its corresponding mIF nodes to encode spatial co-localization.

To effectively capture topological dependencies, we employ a graph attention network^41^ (GAT) to compute adaptive edge weights. For a given node pair (*i,j*), the attention coefficient *e*_*i,j*_ is computed via a shared attentional mechanism:

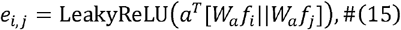

where *f*_*i*_ represents the 256-dimensional input feature of node *i, W*_*a*_ is a learnable weight matrix, *a* is the attention vector, LeakyReLU is the leaky rectified linear unit activation function^72^, and “||” denotes concatenation. These coefficients are normalized using the softmax function to obtain the final attention weights:

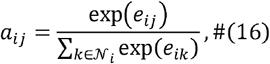

The aggregated contextual feature *z*_*i*_ is then computed as the weighted sum of neighbors, corresponding to the convolution operation:

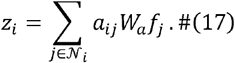

To explicitly model the interaction between the node’s intrinsic features and its gathered context, we introduce a multiplicative update rule. The final node update combines the aggregated context *z*_*i*_ and the projected intrinsic feature *h*_*i*_ through a learnable gate *β*:

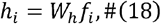

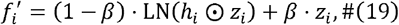

where “⊙” denotes element-wise multiplication, LN applies Layer Normalization^73^, and *β* is a sigmoid-activated learnable scalar. This formulation allows the model to adaptively balance between the linear context aggregation *z*_i_ and the high-order polynomial interaction term *h*_i_ . Note that both and share the same hidden dimension (*d*=256), ensuring that the resulting output 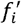 maintains the dimensionality of the feature space. Standard graph attention networks^41^ typically rely on additive aggregation, which may act as a low-pass filter and smooth distinct features. By introducing the multiplicative term *h*_i_ ⊙ *z*_i_, we enable the model to perform feature modulation, where the intrinsic features can selectively amplify or suppress specific channels in the contextual information.

Unlike opaque models, our architecture naturally provides interpretability through the attention mechanism. The attention weight explicitly quantifies the importance of neighbor *j* to node *i* in a patch. By extracting these weights, we can spatially visualize which immune markers drive the model’s decision-making.

#### 4.3 Global node feature updating via graph transformer

Following the integration of mIF information into H&E nodes by polynomial node feature optimization, we extend the H&E graph framework to update node representations from a global perspective. Specifically, we employ a Graph Transformer module to model long-range dependencies among all nodes, enabling the network to capture patterns that span across the whole-slide image. Given the node features matrix *F* within the 256-dimensional space, the graph transformer^50^ updates the node representations as follows:

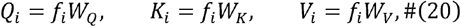

where *W*_*Q*_, *W*_*K*_, and *W*_*V*_ denote the learnable projection matrices. The global attention coefficients are computed via the scaled dot-product:

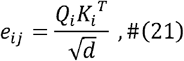

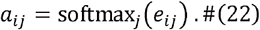

The self-attention output 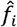 is computed as the weighted sum of values:

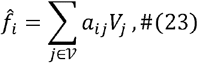

where 𝒱 represents all the nodes. Following the attention mechanism, we employ residual connections^61^ and Layer Normalization^73^ (LN) to stabilize training. The features are then processed by a feed-forward network^66^ (FFN), which projects features to a higher dimension (4*d* = 1024) and then back to *d* (256). The final updated representation 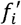 is formulated as:

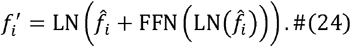

This mechanism extracts structure-aware global features, effectively modeling patterns that span the entire tissue slide.

#### 4.4. Disease risk and development pattern analysis

After polynomial context modeling and global node feature update, we obtain the refined representations for all nodes in the WSI. To derive a patient-level representation, we apply a global average pooling (GAP) function over all node features, resulting in a 256-dimensional compact graph embedding *F*_i_ for patient *i*. Subsequently, we employ a linear projection layer with four output nodes to predict the probability distribution over discrete survival-risk groups. Specifically, the continuous survival time of each sample is discretized into four intervals corresponding to different risk levels.

To optimize the model, we adopt the negative log-likelihood^74^ (NLL) loss in the discrete-time survival setting for patient is defined as:

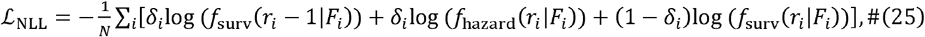

where *F*_*i*_ denotes the features of patient *i. r*_*i*_ is the ground truth time interval, and δ_*i*_ is the censoring indicator (*δ*_*i*_ = 0 indicates censoring, while *δ*_*i*_ =1indicates an observed event). Here, *f*_hazard_ (*r*_*i*_ | *F*_*i*_) represents the conditional hazard probability of experiencing the event at interval *r*_*i*_, given survival up to interval *r*_i_ −1. The survival probability *f*_surv_ (*r*_*i*_ − 1| *F*_*i*_) is explicitly defined as:

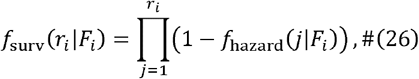

which denotes the probability of surviving through all intervals up to *r*_*i*_.

To quantify the patient-level risk during inference, we derive a scalar risk score from the predicted survival probabilities. Since the summation of survival probabilities across all intervals approximates the expected survival time, we define the risk score *R*_*i*_ as the negative sum of the cumulative survival curve:

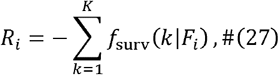

where *K* represents the total number of time intervals. Under this formulation, lower cumulative survival probability results in a higher HCCExplorer risk score, indicating a higher hazard.

The number of time intervals *K* was set to 4 in this work. The training of the survival prediction model is conducted on 4 NVIDIA GeForce RTX 4090 24G GPUs, with a batch size of 1 per GPU using the Adam optimizer of a learning rate of 0.0005. For evaluation, we report the concordance index^75^ (C-index), which measures the proportion of all comparable patient pairs whose predicted risk ordering is consistent with the observed survival times. A higher C-index indicates superior agreement between predicted survival risk and actual clinical outcomes.

### 5 Immune feature extraction in mIF images

To quantitatively characterize the tumor immune microenvironment (TIME) within each 256 × 256 patch, we performed systematic feature extraction using a custom computational pipeline^76,77^ (Figure 4a). This process transformed raw mIF images into 244 high-dimensional, interpretable spatial metrics, where the features were documented in Supplementary Information Table S3.

#### 5.1 Cell detection and phenotyping

The feature extraction pipeline began with precise single-cell segmentation and classification: (i) Cell segmentation: Individual cells were detected and segmented based on the DAPI channel using StarDist^59^, a deep-learning-based method optimized for star-convex polygons to handle dense cellular regions. (ii) Cell identification: Based on the expression levels of specific protein markers (CD3, CD4, CD8, CD19, CD68, and Foxp3), cells were classified into discrete lineages, including CD4+ T cells, CD8+ T cells, Tregs, double-negative (DN) T cells, macrophages, and B cells^54^. (iii) Morphological profiling: For every detected cell, we recorded geometric properties such as nuclear area and density proxies (Supplementary Information Table S3).

#### 5.2 Patch-level feature engineering

Following cell-level identification, we aggregated data to generate patch-level descriptors across three primary domains:

##### 5.2.1 Compositional and statistical features

We calculated the density and relative proportions (fractions) of each cell type within the patch (e.g., frac_CD8T, type_frac B cell). Statistical distributions of marker intensities were captured using metrics such as mean, median, skewness, and kurtosis to reflect the protein expression landscape (e.g., CD3 skew, CD19 kurt). We also computed biological ratios of clinical interest, such as the CD8+ T cell-to-suppressive cell ratio (CD8_to_suppressive_ratio) and the T-to-B cell ratio (T_B_ratio).

##### 5.2.2 Spatial network and graph topology

To model the spatial organization of the TIME, we constructed a patch-level graph where each cell served as a node. (i) Graph construction: Edges were established based on spatial proximity (using a radius or K-nearest neighbor approach), with weights often assigned based on the inverse distance between cell centroids. (ii) Connectivity metrics: We calculated global graph properties, including edge density, the number of connected components, and the diameter of the largest connected component (LCC). (iii) Centrality measures: To identify influential “hub” cells or critical biological “bridges,” we computed degree, betweenness, closeness, and eigenvector centralities (e.g., bet_mean, eig_median). (iv) Community structure: The Louvain algorithm^78^ was employed to detect modular cellular communities, with the resulting modularity and community entropy scores reflecting the degree of spatial clustering.

##### 5.2.3 Cellular interaction and neighborhood analysis

We quantified the spatial relationships between different cell types and marker expressions: (i) Type assortativity: The assort type coefficient was used to measure whether cells of the same lineage tended to aggregate or intermingle with other types. (ii) Edge fractions: We calculated the proportion of edges connecting specific cell types (e.g., edge_frac_T_B) or cells expressing specific markers (e.g., edge_frac_CD3_pos_CD8_pos), providing a direct measure of spatial co-occurrence and potential cellular crosstalk.

All extracted features, including their statistical definitions and biological interpretations, are detailed in Table S3 of the Supplementary Information.

### 6 Immune feature analyses

To interpret the biological decision-making process of HCCExplorer and identify specific components of the tumor immune microenvironment that drive prognostic predictions, we performed a comprehensive post hoc analysis linking deep learning-derived attention weights to quantitative immune features derived from the generated virtual mIF data (Figure 4a).

#### 6.1 Correlation between immune features and attention

First, we elucidated the spatial focus of the model by extracting attention weights from the graph transformer module of HCCExplorer. Since the graph construction is based on super-patches, individual constituent patches (256 × 256 pixels) within a super-patch share the same attention score. These attention values were min-max normalized to the [0, 1] range for each WSI to ensure comparability. To characterize the morphological and molecular phenotypes prioritized by the model, we utilized 110 samples from the internal testing set (65 high-risk and 45 low-risk patients) and stratified tissue patches into high-attention (top 10%) and low-attention (bottom 10%) categories. We then performed a univariate analysis to calculate the correlation between these attention strata and a diverse set of quantitative patch-level immune features (e.g., cell counts, expression intensity, spatial entropy) obtained from the virtual mIF workflow. To account for inter-patient variability, immune features were standardized via group-level normalization. Crucially, this correlation analysis was conducted independently in the low- and high-risk patient cohorts to disentangle the specific immune signatures that drive the model’s risk stratification across different prognostic contexts (Figure 4b, c).

#### 6.2 Feature quantification and survival analysis at the slide-level

Building on the correlation analysis, we selected the 34 immune features with the strongest associations with the model’s attention mechanism across risk groups. To evaluate the clinical prognostic value of these biologically interpretable features, we aggregated patch-level metrics into slide-level representations by computing the mean of each feature across all valid tissue patches in a WSI. We subsequently performed univariate Cox proportional hazards regression analyses^79^ to assess the association of these aggregated immune features with clinical outcomes on the FAZJU cohort, specifically overall survival and disease-free survival. For each feature, patients were stratified into high- and low-expression groups based on optimal cutoff values determined by the maximally selected rank statistics, and Kaplan-Meier curves^80^ were generated to assess the statistical significance of the prognostic separation. To ensure rigorous validation, these specific cutoff thresholds derived from the training set were applied unchanged to stratify patients across all independent external validation cohorts (Figure 4d, e). Kaplan-Meier curves were then generated to visualize the statistical significance of the prognostic separation across all datasets (Figure 4g, Extended Data Figure 6).

#### 6.3 Immune feature visualization

To provide an intuitive spatial interpretation of the heterogeneous immune landscape, we generated high-resolution whole-slide heatmaps for key immune features (Figure 4f, Extended Data Figure 6). We mapped the quantitative values of selected immune markers (e.g., macrophage density) directly back to their original spatial coordinates on the WSI grid. To ensure visual clarity and focus strictly on valid tissue regions, patches containing undefined values (N/A) or background noise were rendered transparent. This visualization strategy facilitated a direct visual comparison of H&E morphology, the generated virtual mIF signals, and the derived quantitative immune maps, thereby validating the spatial consistency of the model’s biological insights.

#### 6.4 Immune feature spatial analyses

Through the graph transformer^50^, we extracted attention weights assigned to the edges linking source and target nodes (Figure 6a, b, Extended Data Figures 7, 8). Each super-patch is represented by its central patch with 256 × 256 pixels, labeled according to its majority cell population. By aggregating these weights, we derived quantitative measures of cross-region and cell-type interactions, providing interpretable spatial insights within the HCCExplorer framework. In addition to the frequency statistics shown in Figure 6d, the attention weight statistics were presented in Supplementary Information Figure S1. The core conclusions largely aligned with those derived from the frequency statistics.

### 7 Cell-type deconvolutions of bulk RNA-seq

To validate the findings from our virtual mIF pipeline at a transcriptomic level and further refine cellular subtypes, we performed computational deconvolution on bulk RNA sequencing (RNA-seq) data.

#### 7.1 Data acquisition and processing

We utilized the TCGA-LIHC^55^ dataset, which provides comprehensive bulk RNA-seq profiles from patients with HCC. The raw expression data were normalized and transformed into a format compatible with deconvolution algorithms to ensure consistency across samples.

#### 7.2 CIBERSORTx deconvolution

The cellular composition of the HCC bulk tissues was estimated using CIBERSORTx^45^, a machine learning framework that infers cell-type-specific gene expression profiles and proportions from bulk tissue. (i) Signature matrix: We employed a validated signature matrix (LM22^81^) to guide the deconvolution process. (ii) Algorithm execution: The deconvolution was run with batch correction and 100 permutations to ensure the statistical significance of the estimated cell fractions. More detailed descriptions are in https://cibersortx.stanford.edu/.

#### 7.3 Integration of virtual mIF-derived counts and subtype refinement

A key strength of our methodology lies in the integration of spatial imaging data with transcriptomic deconvolution: (i) Macrophage quantification: While our virtual mIF pipeline provided the absolute counts and spatial distribution of total macrophages (CD68+ cells) within tissue patches, CIBERSORTx allowed us to dissect the underlying functional heterogeneity of these cells. (ii) Subtype inference: By applying the proportions of macrophage subtypes (e.g., M0, M1, and M2) derived from CIBERSORTx to the total macrophage counts obtained via mIF, we were able to estimate the absolute abundance of specific macrophage functional states within the tumor microenvironment.

### 8 Evaluation metrics

#### 8.1 Evaluation of H&E-to-virtual mIF translation

To comprehensively assess the quality of the generated virtual mIF images, we employed a multifaceted strategy that combined quantitative computational metrics (FID, CSS) and qualitative expert assessment (visual Turing test, staining quality) on HCCExplorer and six state-of-the-art image-to-image translation methods (Supplementary Information Table S1):

i. **CycleGAN**^36^: The foundational framework for unpaired image-to-image translation that utilizes a cycle-consistency loss to enforce structural alignment between source and target domains. We trained CycleGAN on the same training sets with the HCCExplorer to obtain the final weights for evaluation.
ii. **CUT**^34^: CUT introduces patch-wise contrastive learning (PatchNCE) to maximize mutual information between corresponding patches, enabling effective one-sided translation without the need for cycle consistency. As the baseline of C3UT, we trained the CUT on the same training sets with the HCCExplorer to obtain the final weights for evaluation.
iii. **FastCUT**^34^: FastCUT is a lightweight and computationally efficient variant of CUT designed for faster training and reduced memory overhead while maintaining high-quality translation performance. We trained the FastCUT on the same training sets with the HCCExplorer to obtain the final weights for evaluation.
iv. **AI-FFPE**^35^: A specialized medical imaging model that uses generative adversarial networks to virtually transform frozen section pathology slides into high-quality, permanent-section (FFPE) equivalents. We trained the AI-FFPE on the same training sets with the HCCExplorer to obtain the final weights for evaluation.
v. **VirtualMultiplxer**^24^: A deep learning approach that synthesizes virtual multiplexed immunohistochemistry (mIHC) images from standard H&E-stained slides. We trained the VirtualMultiplxer on the same training sets with the HCCExplorer to obtain the final weights for evaluation.
vi. **GigaTIME**^25^: GigaTIME is a multi-modal AI framework for population-scale TIME modeling by bridging cell morphology and states. GigaTIME learns a cross-modal translator to generate virtual mIF images from hematoxylin and eosin (H&E) slides by training on 40 million cells with paired H&E and mIF data across 21 proteins. Here, we directly used the trained GigaTIME on the H&E slides of HCC for evaluation.

##### 8.11 Fréchet Inception Distance (FID)

To evaluate the visual realism and distribution consistency of the generated mIF patches compared to real mIF patches, we utilized the Fréchet Inception Distance^37^ (FID). FID measures the distance between the feature distributions of real (*r*) and generated (*g*) images in the deep feature space of a pre-trained Inception-v3^82^ network.

Lower FID scores indicate that the generated images are more similar to the real data in terms of visual quality and diversity. The metric is defined as:

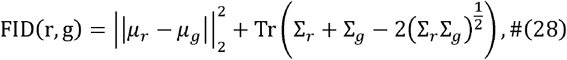

where μ_*r*_ and μ_*g*_ represent the mean feature vectors of the real and generated distributions, respectively. ∑_*r*_ and ∑_*g*_ denote their corresponding covariance matrices. Tr refers to the trace of the matrix. This metric is particularly effective at detecting mode collapse and artifacts in generative models.

##### 8.1.2 Contrast-Structure Similarity (CSS)

While FID assesses distribution-level realism, it is crucial to verify that the generated images preserve tissue’s the fine-grained histological architecture and signal intensity. To this end, we employed the Contrast-Structure Similarity^38^ (CSS) index. CSS is derived from the Structural Similarity Index Measure^83^ (SSIM) but specifically emphasizes the preservation of image contrast and structures. For two image patches *r* (real) and *g* (generated), the CSS index is calculated as:

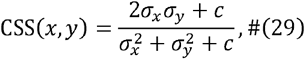

where *σ* _*x*_ and *σ* _*y*_ are the standard deviations, and is the stability constant.

##### 8.1.3 Visual Turing test

To quantify the perceptual indistinguishability of the generated virtual mIF images from real mIF images, we conducted a rigorous visual Turing test involving four board-certified pathologists, each with over five years of experience. For each marker, we curated a blinded evaluation dataset consisting of a balanced set of 50 H&E-real mIF pairs and 50 H&E-virtual mIF pairs. The pathologists were blinded to the origin of the images and were independently tasked with classifying each patch as either “Real” or “Virtual.” The classification accuracy, sensitivity, and specificity were calculated to determine if the generated images could successfully deceive human experts (where a classification accuracy close to 50% indicates true indistinguishability).

##### 8.1.4 Staining quality assessment

Beyond perceptual realism, we further assessed the biological fidelity and clinical utility of the virtual staining. The same cohort of four pathologists performed a comprehensive qualitative assessment on a set of 50 real and 50 virtual images per marker. The staining quality was rigorously evaluated based on five key histological criteria derived from clinical standards: (i) overall expression intensity, (ii) signal-to-noise ratio (background noise), (iii) consistency of staining patterns, (iv) cell-type specificity, and (v) accuracy of subcellular localization (e.g., distinguishing between nuclear and membranous signals). Pathologists were asked to grade the staining quality of the virtual images as clinically acceptable and comparable to the standard of care for real mIF staining.

#### 8.2 Benchmarking survival prediction methods

To validate the superior prognostic performance of HCCExplorer, we benchmarked our HCCExplorer against clinical standards and seven distinct survival analysis methods, representing a broad spectrum of established baselines and state-of-the-art foundation models (Supplementary Information Table S2). The comparison methods included:

(i) **UNI-h2**^29^ **+ average pooling (Avg-Pool)**: UNI^29^ is a general-purpose self-supervised model for pathology, pretrained using more than 100□million images from over 100,000 diagnostic H&E-stained WSIs. The UNI-h2^29^ is an upgraded version of UNI, which trained on over 200 million pathology H&E and IHC images sampled from 350+ thousand diverse WSIs. Here, we used UNI-h2 as the H&E feature extractor to obtain patch embeddings, and use the average pooling to aggregate patch embeddings into the bag feature for survival prediction.
(ii) **UNI-h2**^29^ **+ ABMIL**^40^: We used UNI-h2 for H&E patch feature encoding, and used a gated attention mechanism to weigh patch embeddings.
(iii) **UNI-h2**^29^ **+ GAT**^41^: We used UNI-h2 for H&E patch feature encoding, and used a graph attention mechanism to aggregate patch embeddings.
(iv) **CHIEF**^30^: Clinical Histopathology Imaging Evaluation Foundation (CHIEF) model is a general-purpose weakly supervised machine learning framework to extract pathology imaging features for systematic cancer evaluation. CHIEF leveraged two types of pretraining procedure: unsupervised pretraining on 15 million unlabelled tile images and weakly supervised pretraining on more than 60,000 WSIs. Here, we used the pre-trained CHIEF as the H&E WSI feature encoder, and fine-tuned the WSI feature for survival prediction.
(v) **Prov-GigaPath**^31^: Prov-GigaPath is a whole-slide pathology foundation model pretrained on 1.3 billion256 × 256 pathology image tiles in 171,189 whole slides from Providence, a large US health network comprising 28 cancer centres. Here, we used the pre-trained Prov-GigaPath as the WSI feature encoder, and fine-tuned the H&E WSI feature for survival prediction.
(vi) **TITAN**^32^: Transformer-based pathology Image and Text Alignment Network (TITAN) is a multi-modal whole-slide foundation model pretrained using 335,645 whole-slide images via visual self-supervised learning and vision-language alignment with corresponding pathology reports and 423,122 synthetic captions generated from a multi-modal generative AI copilot for pathology. As the upgraded version of CONCH^84^, we used the pre-trained TITAN as the WSI feature encoder, and fine-tuned the H&E WSI feature for survival prediction.
(vii) **KRONOS**^33^: KRONOS is a foundation model built for spatial proteomics, which was trained in a self-supervised manner on over 47 million image patches covering 175 protein markers, 16 tissue types, and 8 fluorescence-based imaging platforms. Here, we used the pre-trained KRONOS as the mIF feature encoder for obtaining patch-level features, and adopt the average pooling to obtain the mIF features for survival prediction.

##### 8.2.1 Generalization evaluation protocol

A pivotal objective of this study was to comprehensively assess the cross-center generalization capability of the developed models. To achieve this, we adhered to a strict train-once, test-everywhere protocol, where all models were trained exclusively on the OS data from the FAZJU internal training set. Crucially, the training phase relied solely on OS supervision, with no information regarding DFS and RFS incorporated. These OS-trained models were subsequently deployed for direct inference across the FAZJU internal test set and three independent external validation cohorts (SAZJU, TCGA, and YWCH). In this phase, the risk scores derived purely from OS learning were utilized to stratify patients for all survival endpoints, including DFS and RFS, without any re-training, fine-tuning, or domain adaptation. This rigorous experimental setting provides a robust evaluation of model performance across diverse data sources and scanning protocols, while simultaneously demonstrating that the extracted risk features capture fundamental tumor aggressiveness transferable across different clinical outcomes.

##### 8.2.2 Performance metrics

We employed a 5-fold cross-validation to evaluate the model performance. The mean C-index^75^ and its standard deviation (SD) were calculated across the five folds. The statistical comparisons between models were performed using a two-sided paired t-test on the results of the 5 cross-validation folds. To evaluate risk stratification, Kaplan-Meier (KM) curves^85^ and the Log-rank test^86^ were employed. Furthermore, Cox proportional hazards regression^87^ was performed to estimate Hazard Ratios (HRs) and their associated *P* values (Wald test^88^), quantifying the contribution of the extracted features to predicted survival risk.

## Data Availability Statement

For benchmarks, TCGA-LIHC data can be accessed through the NIH Genomic Data Commons (https://portal.gdc.cancer.gov). In accordance with institutional policies, all requests for data collected or curated in-house will be evaluated on a case-by-case basis to determine whether the requested data complies with intellectual property and patient privacy obligations.

## Code Availability Statement

The source code for HCCExplorer is publicly available for academic research purposes at https://github.com/MedCAI/HCCExplorer. The model weights will be released.

## Acknowledgements

W.L. was funded by the Huadong Medicine Joint Fund of the Zhejiang Provincial Natural Science Foundation of China (grant No. LHDMZ24H160001). Y.Z. was funded by the National Natural Science Foundation of China under 62331011. X.Z. was funded by the Huadong Medicine Joint Fund of the Zhejiang Provincial Natural Science Foundation of China under Grant No. LHDMZ25H160002, the Zhejiang Province Health Major Science and Technology Program of National Health Commission Scientific Research Fund (No. WKJ-ZJ-2426). J.N.K. is supported by the German Cancer Aid DKH (DECADE, 70115166), the German Federal Ministry of Research, Technology and Space BMFTR (PEARL, 01KD2104C; CAMINO, 01EO2101; TRANSFORM LIVER, 031L0312A; TANGERINE, 01KT2302 through ERA-NET Transcan; Come2Data, 16DKZ2044A; DEEP-HCC, 031L0315A; DECIPHER-M, 01KD2420A; NextBIG, 01ZU2402A; PROSURV, 01KD2509C), the German Research Foundation (DFG, Deutsche Forschungsgemeinschaft) as part of Germany’s Excellence Strategy – EXC 2050/2 – Project ID 390696704 – Cluster of Excellence “Centre for Tactile Internet with Human-in-the-Loop” (CeTI) of Technische Universität Dresden, as well as through DFG-funded collaborative research projects (TRR 412/1, 535081457; SFB 1709/1 2025, 533056198), the German Academic Exchange Service DAAD (SECAI, 57616814), the German Federal Joint Committee G-BA (TransplantKI, 01VSF21048), the European Union EU’s Horizon Europe research and innovation programme (ODELIA, 101057091; GENIAL, 101096312), the European Research Council ERC (NADIR, 101114631), the Breast Cancer Research Foundation (BELLADONNA, BCRF-25-225) and the National Institute for Health and Care Research NIHR (Leeds Biomedical Research Centre, NIHR203331). The views expressed are those of the author(s) and not necessarily those of the NHS, the NIHR or the Department of Health and Social Care. This work was funded by the European Union. Views and opinions expressed are, however, those of the author(s) only and do not necessarily reflect those of the European Union. Neither the European Union nor the granting authority can be held responsible for them.

## Author contributions

L.C., S.J., J.L., F.L., B.Z., X.Z., J.Z., J.N.K., Y.Z., and W.L. conceptualized the study and its design. L.C., S.J., J.L., and Y.Z. created the methodology. L.C., F.L., J.Z., N.G., Q.Z., and J.N.K. conducted the investigation. L.C., S.J., F.L., B.Z., Q.Z, Z.H., Y.M., Z.L., S.F., M.H., X.Z, J.Z., and W.L. were involved in data collection and pre-processing. L.C., S.J., J.L., F.L., Y.Z., and W.L. performed the data analysis and visualized the study. X.Z., J.N.K, Y.Z., and W.L. completed the project administration. L.C., S.J., F.L., and B.Z. drafted the manuscript. J.L, J.Z., J.N.K., Y.Z., and W.L. critically reviewed, edited, and approved the manuscript.

## Extended Data

**Extended Data Figure 1.**
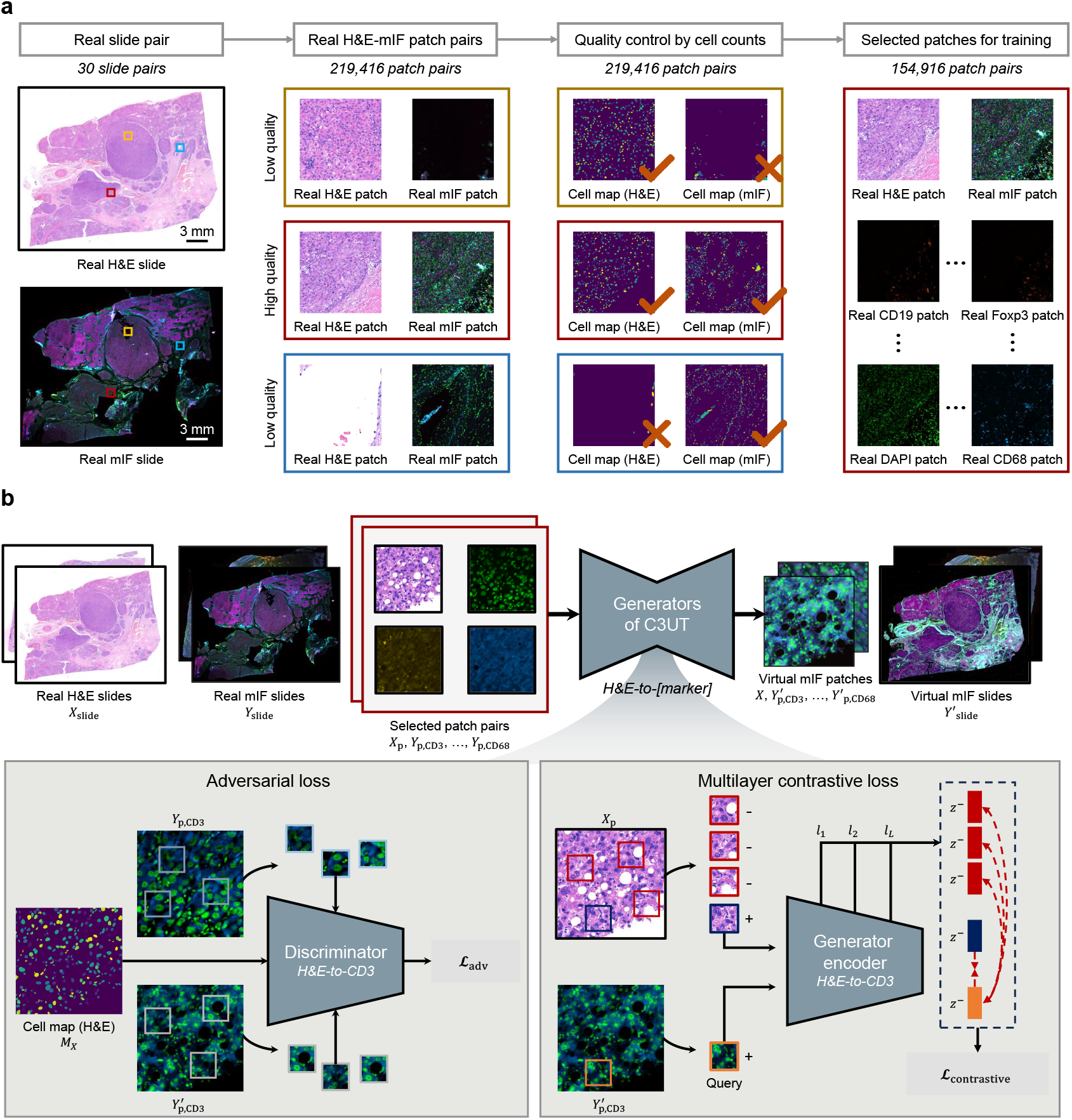
H&E-to-mIF translation dataset construction pipeline and C3UT model architecture. **a** Data preprocessing and quality control workflow. 30 pairs of real H&E and multiplex immunofluorescence (mIF) whole-slide images (WSIs) from consecutive sections were co-registered. A total of 219,416 patch pairs were initially extracted. To ensure informational richness, a rigorous quality control mechanism based on cell counts was applied: patch pairs were discarded if they contained background/noise in the mIF domain (top row, orange box) or lacked tissue content in the H&E domain (bottom row, blue box). Only pairs exhibiting meaningful biological signals in both domains were retained (middle row, red box), resulting in a final curated dataset of 154,916 patch pairs for training. **b** Schematic of the cell-consistent cross-modal unpaired translation (C3UT) framework. The model takes real H&E slides (*X* _s□ide_) as input to generate virtual mIF slides 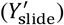 via patch-based processing. Left box (Adversarial loss): Illustration of the cell-map-guided adversarial learning. The discriminator differentiates between real (*Y*_*p*_) and generated 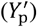 mIF patches while being conditioned on the cell map (*M*_*X*_) derived from the H&E input. This constraint enforces that the generated protein signals spatially align with the underlying nuclear topology. Right box (Multilayer contrastive loss): Illustration of the PatchNCE loss mechanism. Feature representations are extracted from multiple layers (*l*_1_, *l*_2_, …, *l*_*L*_) of the generator encoder. The loss maximizes the mutual information between the generated “query” patch and its spatially corresponding “positive” H&E patch (+), while pushing away “negative” patches (-) sampled from different locations, ensuring semantic correspondence between the histological morphology and the synthesized proteomic signal.

**Extended Data Figure 2.**
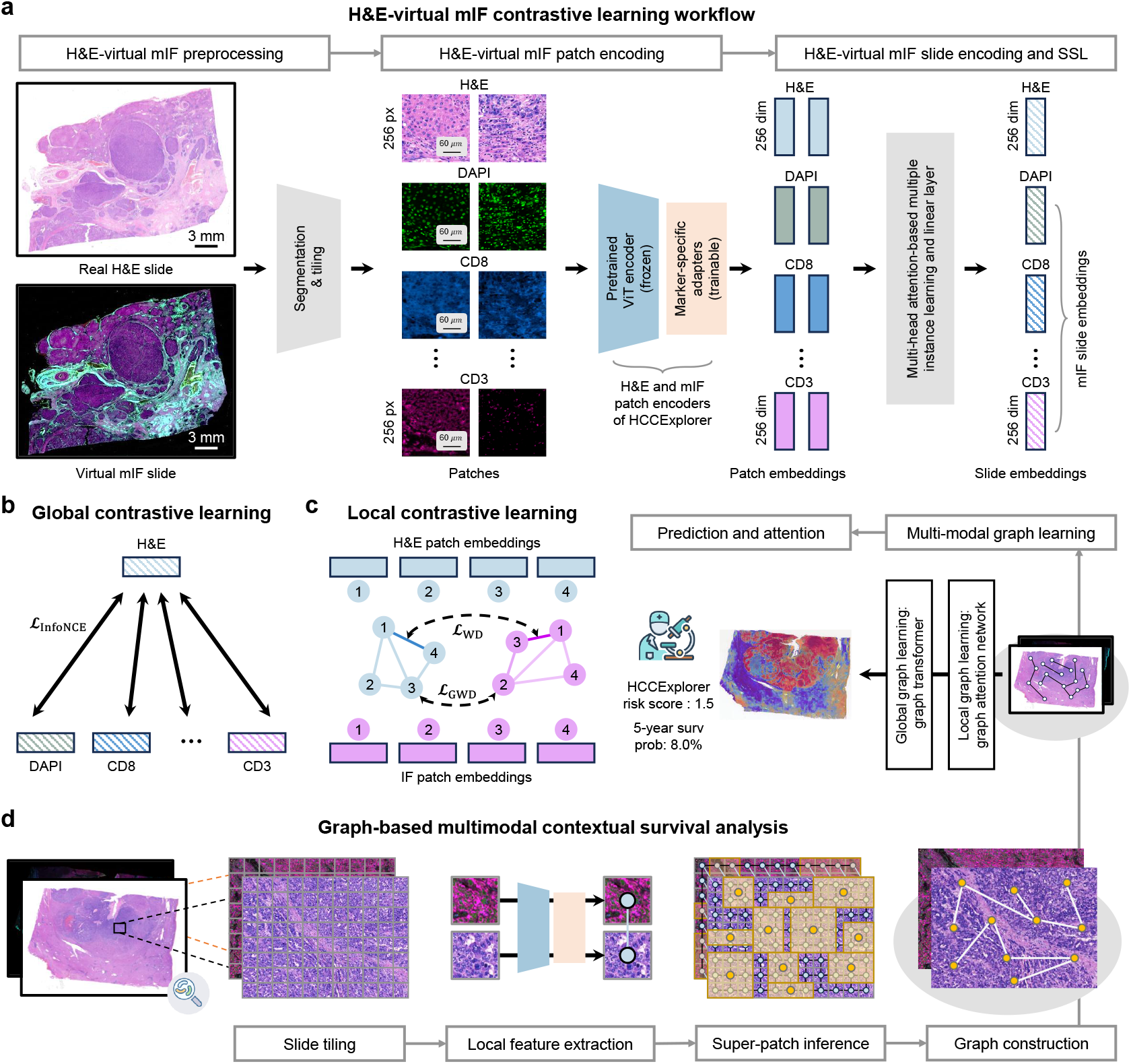
Pipelines of multi-modal contrastive learning and graph-based survival analyses. **a** Workflow for H&E-virtual mIF contrastive learning. Paired real H&E and generated virtual mIF whole-slide images (WSIs) are preprocessed via segmentation and tessellated into non-overlapping 256 × 256 pixel patches. A frozen pre-trained Vision Transformer encoder extracts high-level features, which are subsequently projected into a marker-specific latent space (256 dimensions) via trainable marker-specific adapters (orange boxes). These patch-level embeddings are then aggregated into slide-level representations using a multi-head attention-based multiple instance learning module. **b** Schematic of global contrastive learning. The model minimizes the InfoNCE loss (ℒ _infoNCE_) to align the global slide-level embedding of the H&E modality with the corresponding embeddings of various virtual Mif markers (e.g., CD3 and CD8), ensuring cross-modal consistency at the slide level. **c** Schematic of local contrastive learning and prediction architecture. To enforce fine-grained alignment, a graph optimal transport mechanism is employed, comprising the Wasserstein distance for aligning patch distributions (nodes) and the Gromov-Wasserstein distance for aligning spatial topologies (edges) between H&E and mIF graphs. The right panel illustrates the HCCExplorer architecture, which integrates global graph transformers and local graph attention networks to output a risk score and survival probability. **d** Pipeline for graph-based multi-modal contextual survival analysis. The process involves hierarchical processing, from slide tiling and local feature extraction to super-patch inference, culminating in a spatial graph in which nodes represent tissue regions and edges encode spatial adjacency to model interactions within the tumor microenvironment.

**Extended Data Figure 3.**
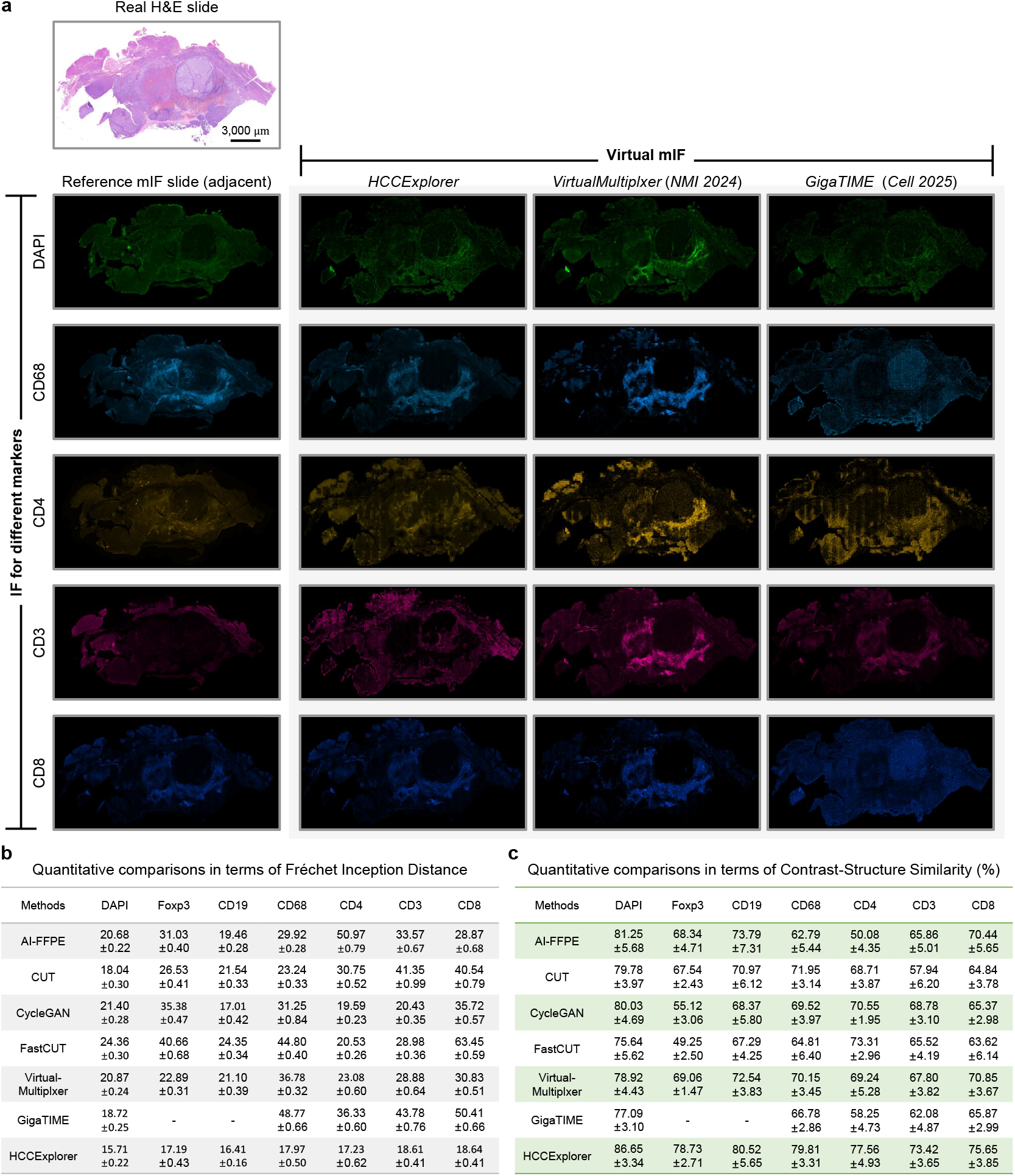
Qualitative and quantitative performance evaluation of H&E-to-virtual mIF translation methods. **a** Qualitative comparison of H&E-to-virtual mIF translation across different markers. A real H&E-stained WSI of HCC is shown as input, alongside an adjacent real mIF slide used as a biological reference. The panels display the synthesized virtual mIF channels (DAPI, CD68, CD4, CD3, and CD8) generated by the proposed HCCExplorer and two advanced models: VirtualMultiplexer (*Nature Machine Intelligence*, 2024) and GigaTIME (*Cell* 2025). HCCExplorer demonstrates superior visual fidelity, capturing the spatial distribution and staining patterns of immune markers that more closely align with the reference mIF slide. **b** Quantitative evaluation using Fréchet Inception Distance (FID). The table summarizes the FID scores for seven different virtual staining methods across seven markers. Lower FID scores indicate higher distributional similarity to real mIF data. HCCExplorer achieves the lowest FID scores across all evaluated markers, outperforming existing generative frameworks. **c** Quantitative evaluation using Contrast-Structure Similarity (CSS) score. The structural integrity of the generated images is assessed via CSS. Higher percentages indicate better preservation of morphological and contrast-based information. HCCExplorer consistently maintains the highest similarity scores across the marker panel, validating its robustness in preserving fine-grained cellular structures during the H&E-to-virtual mIF translation.

**Extended Data Figure 4.**
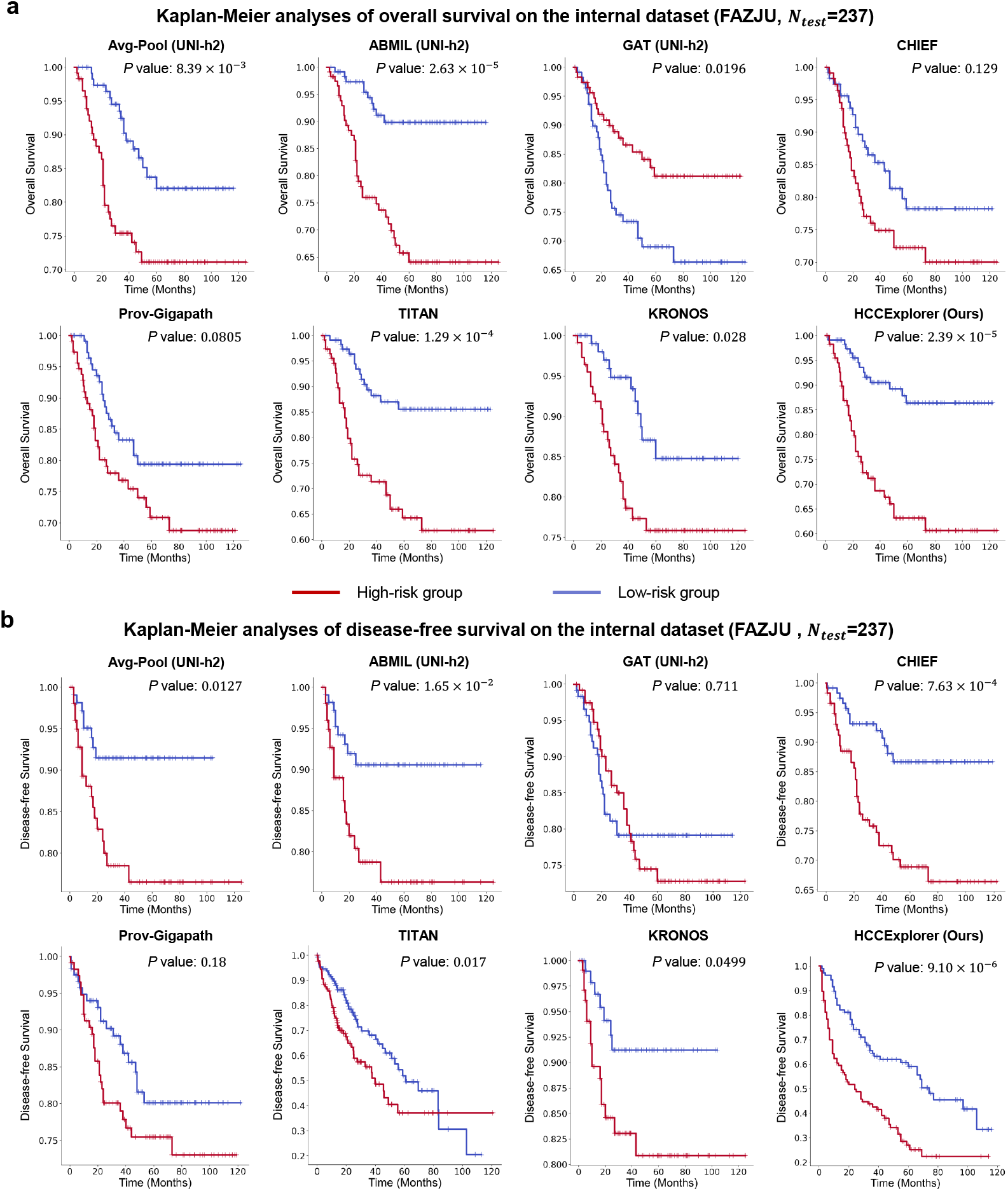
Prognostic performance benchmarking on the internal FAZJU testing set. **a** Kaplan-Meier analysis of overall survival (OS) on the independent internal testing cohort (FAZJU). Patients were stratified into high-risk (red) and low-risk (blue) groups based on the median risk score predicted by each model. HCCExplorer demonstrates superior risk stratification capability (P = 2.39 × 10^−5^, Log-rank test) compared to seven state-of-the-art baseline methods, including Avg-Pool, ABMIL, GAT, CHIEF, Prov-Gigapath, TITAN, and KRONOS. **b** Kaplan-Meier analysis of disease-free survival (DFS) on the same internal testing cohort (FAZJU). HCCExplorer achieves the most significant separation between risk groups (*P* = 9.10 × 10^−6^, Log-rank test), consistently outperforming other competing computational pathology algorithms in predicting tumor recurrence.

**Extended Data Figure 5.**
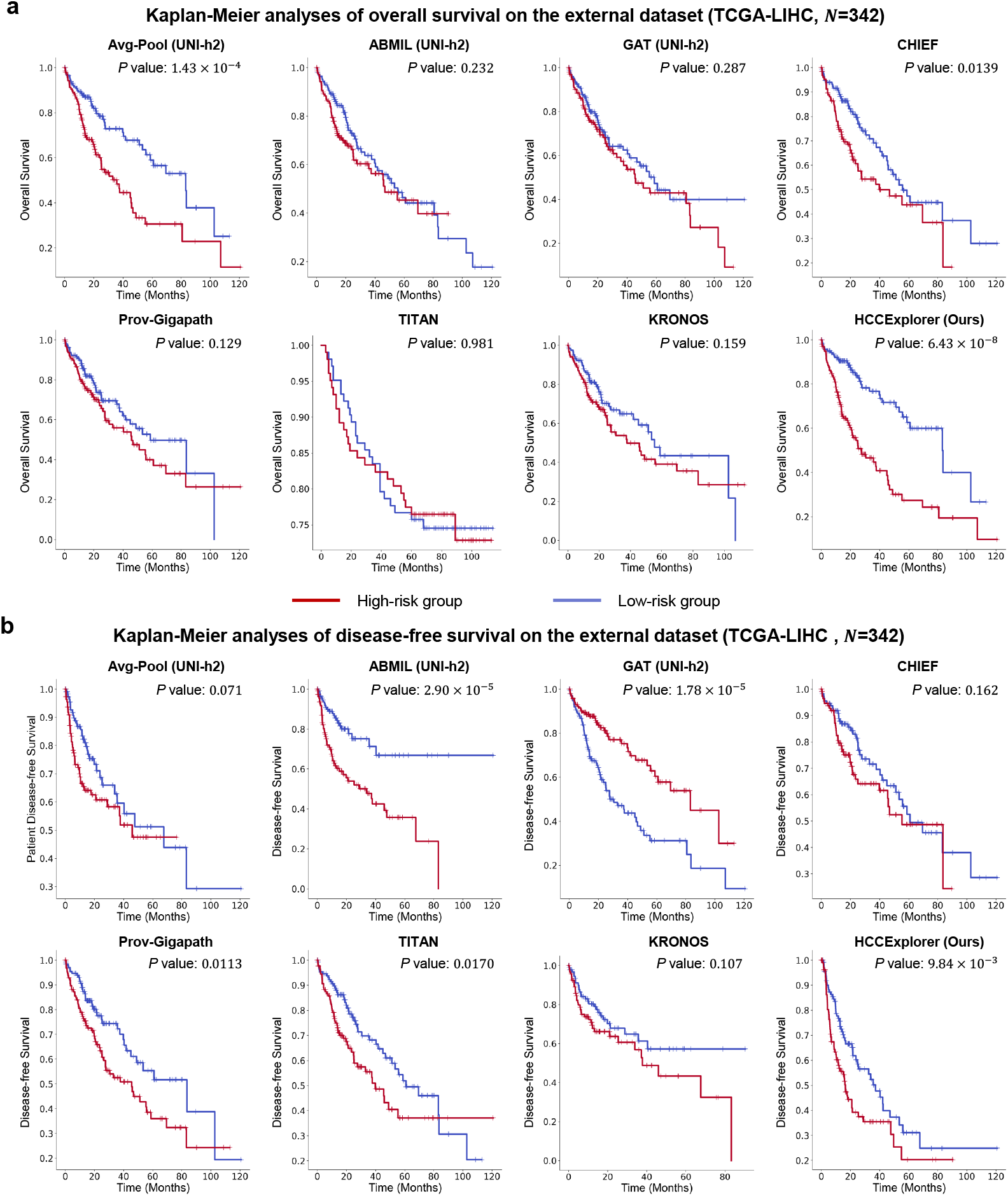
Prognostic performance benchmarking on the external TCGA-LIHC validation set. **a** Kaplan-Meier analysis of overall survival (OS) on the independent external testing cohort (TCGA-LIHC). Patients were stratified into high-risk (red) and low-risk (blue) groups based on the risk scores predicted by each model. HCCExplorer demonstrates robust generalization capability, achieving the most significant risk stratification (*P* =6.43 × 10^-8^, Log-rank test) compared to seven state-of-the-art baseline methods, including Avg-Pool, ABMIL, GAT, CHIEF, Prov-Gigapath, TITAN, and KRONOS. **b** Kaplan-Meier analysis of disease-free survival (DFS) on the external TCGA-LIHC cohort. HCCExplorer successfully stratifies patients into distinct high- and low-risk groups with statistical significance (*P*= 9.84 × 10^−3^, Log-rank test), further validating the model’s prognostic value and transferability across diverse patient populations.

**Extended Data Figure 6.**
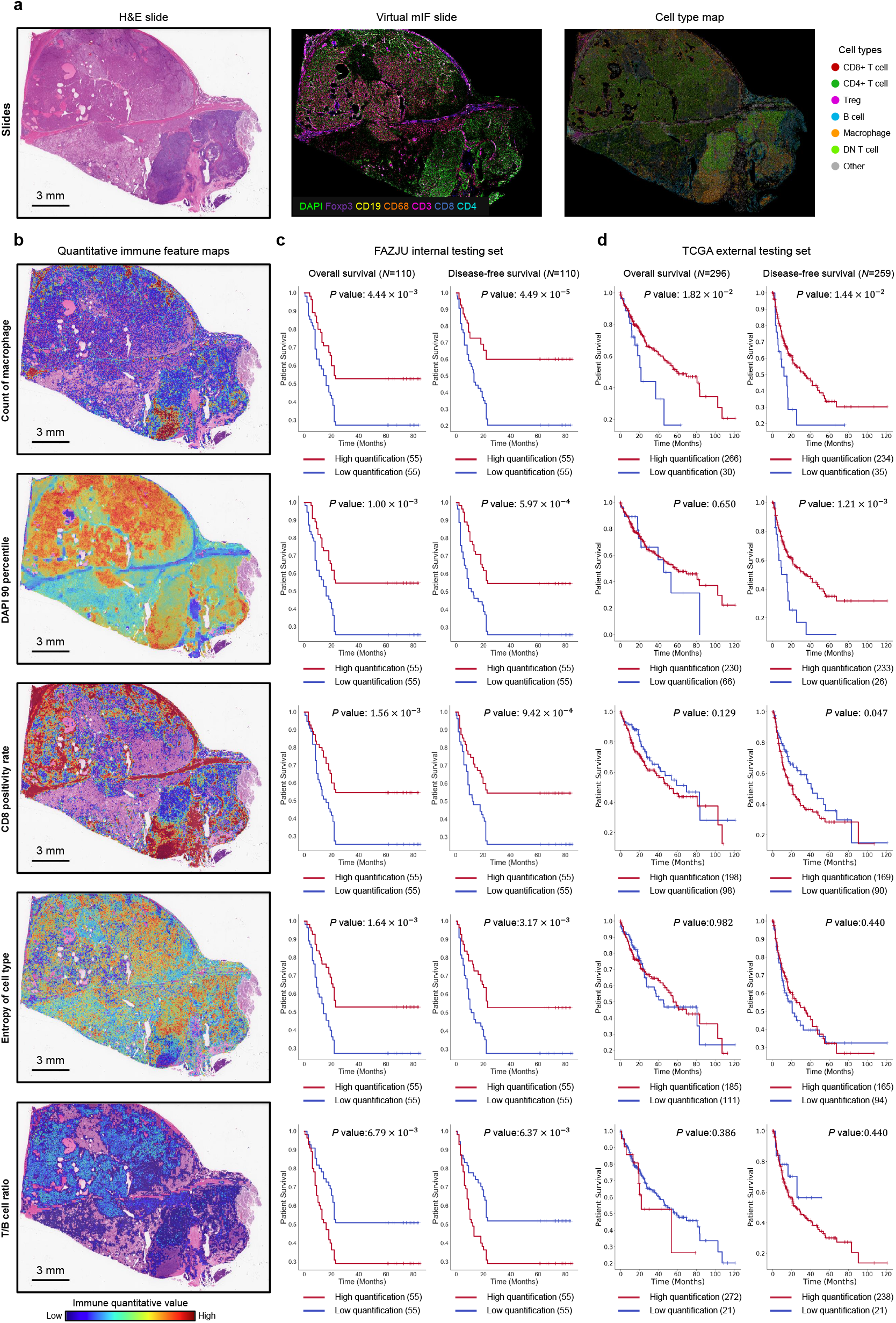
Visualization of spatial quantitative immune features and their prognostic significance. **a** Visualization of the H&E-to-virtual mIF translation for a representative HCC case. **b** Whole-slide spatial heatmaps quantifying five key immune features per patch identified by HCCExplorer. From top to bottom: Count of macrophage, DAPI 90th percentile intensity (reflecting nuclear activity), CD8 positivity rate, Entropy of cell type (representing immune heterogeneity), and T/B cell ratio. The color scale ranges from low (blue) to high (red) feature values, highlighting the spatial heterogeneity of the tumor microenvironment. **c** Kaplan-Meier survival analyses of the five corresponding immune features in the FAZJU internal testing set. Patients were stratified into high (red) and low (blue) groups based on the slide-level quantification of each feature. P values for overall survival (OS) and disease-free survival (DFS) were calculated using the Log-rank test. **d** Independent validation of the prognostic value of the same five immune features in the external TCGA-LIHC testing set. The consistent stratification across cohorts demonstrates the robustness and generalizability of the virtual mIF-derived biomarkers. Note: Immune feature extraction failed for a subset of slides due to computational issues. Consequently, the TCGA cohort shown in the figure comprises only patients with complete feature sets.

**Extended Data Figure 7.**
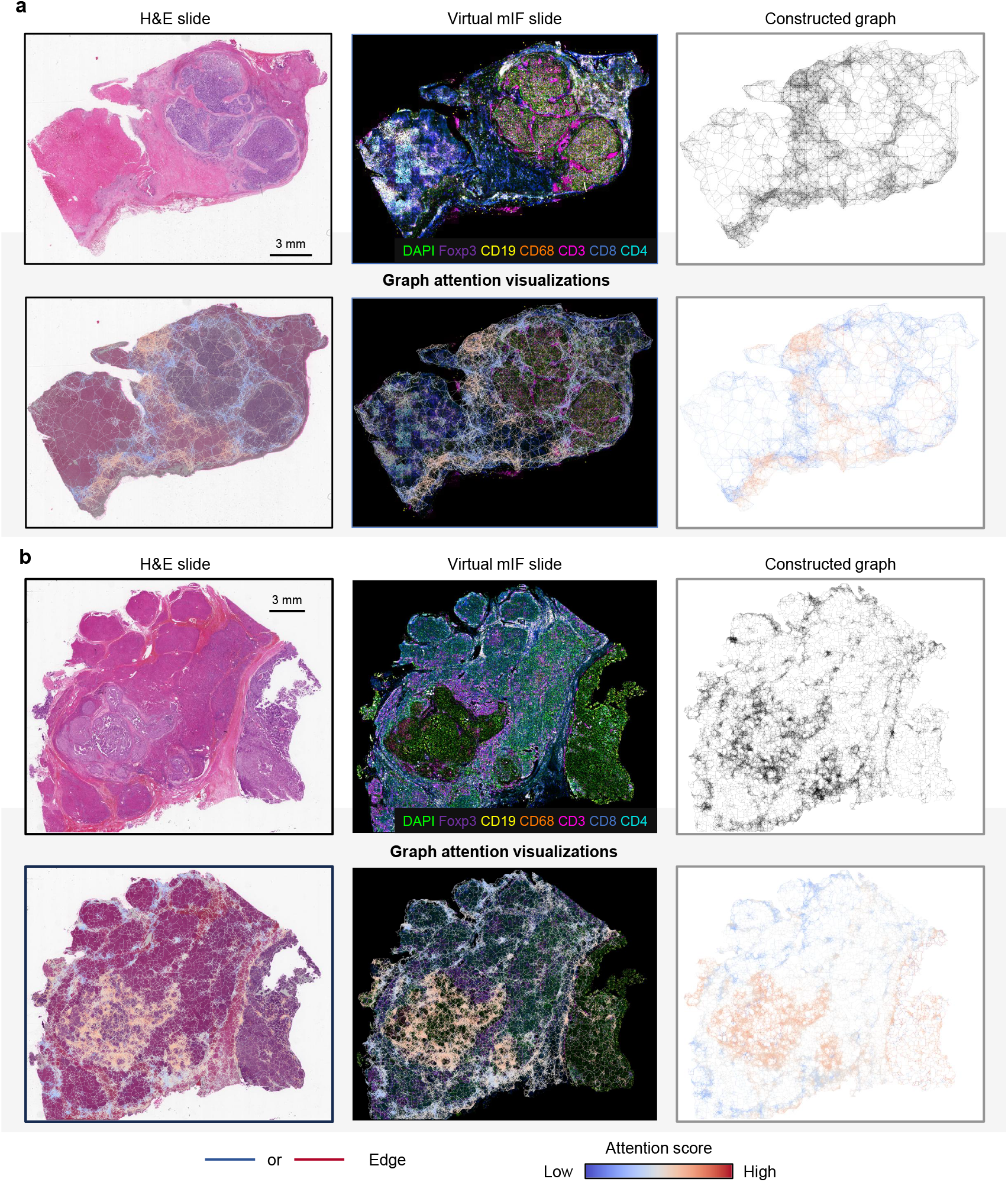
Visualization of multi-modal graph construction and interpretability via multi-modal contextual learning. **a, b** Representative examples of the graph-based modeling pipeline for two independent HCC patients. Top row: Displays the input H&E-stained whole-slide image, the corresponding virtual mIF image, and the constructed spatial graph. In the graph structure, nodes represent aggregated tissue super-patches, and edges represent the spatial connectivity between adjacent regions. Bottom row: Visualizes the graph attention maps derived from the HCCExplorer model. The learned attention weights are back-projected onto the H&E morphology, the virtual mIF landscape, and the graph topology to highlight regions contributing most significantly to the prognostic prediction. The color scale (bottom right) indicates the attention score, where red denotes high attention (regions of high prognostic relevance) and blue denotes low attention. The edges are similarly color-coded to visualize the importance of spatial interactions between nodes.

**Extended Data Figure 8.**
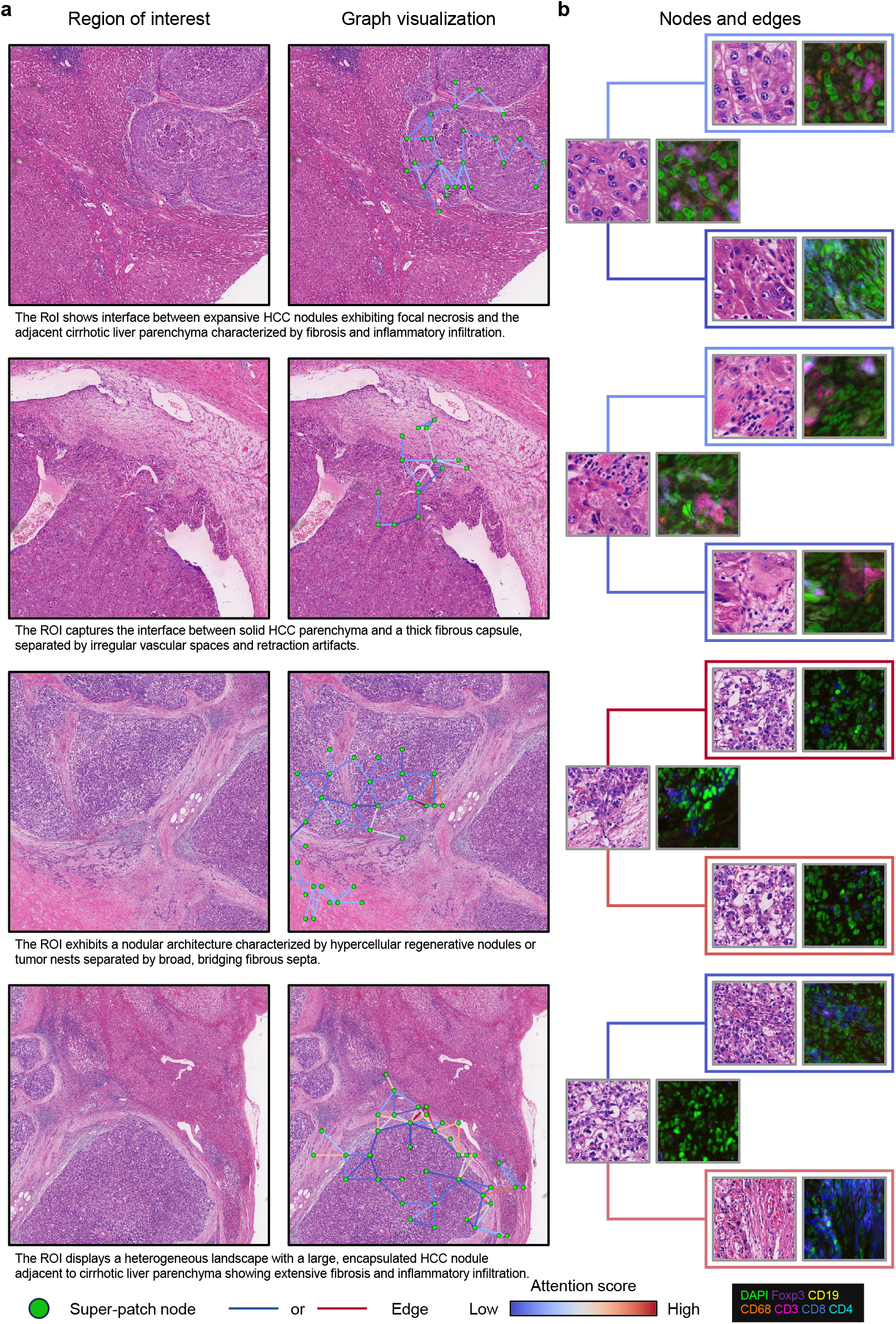
Fine-grained visualization of local spatial graphs and multi-modal interactions within Regions of Interest (RoIs). **a** Visualization of local spatial graph construction across representative histological patterns. The panel displays four distinct RoIs capturing critical tumor immune microenvironment components (from top to bottom): (i) interface between expansive HCC nodules and cirrhotic liver parenchyma; (ii) interface between solid HCC parenchyma and fibrous capsules; (iii) nodular architecture separated by bridging fibrous septa; and (iv) heterogeneous landscape with encapsulated nodules. The left column shows the original H&E staining, while the right column overlays the constructed graph, where green nodes represent tissue super-patches and connecting lines represent spatial edges. **b** Detailed visualization of multi-modal node-edge interactions. Selected pairs of interacting super-patches are magnified to showcase the learned multi-modal representations, displaying both H&E morphology and the corresponding generated virtual mIF signals. The connecting edges are color-coded based on the attention score (ranging from blue/low to red/high), highlighting the model’s ability to prioritize specific spatial interactions.

**Extended Data Figure 9.**
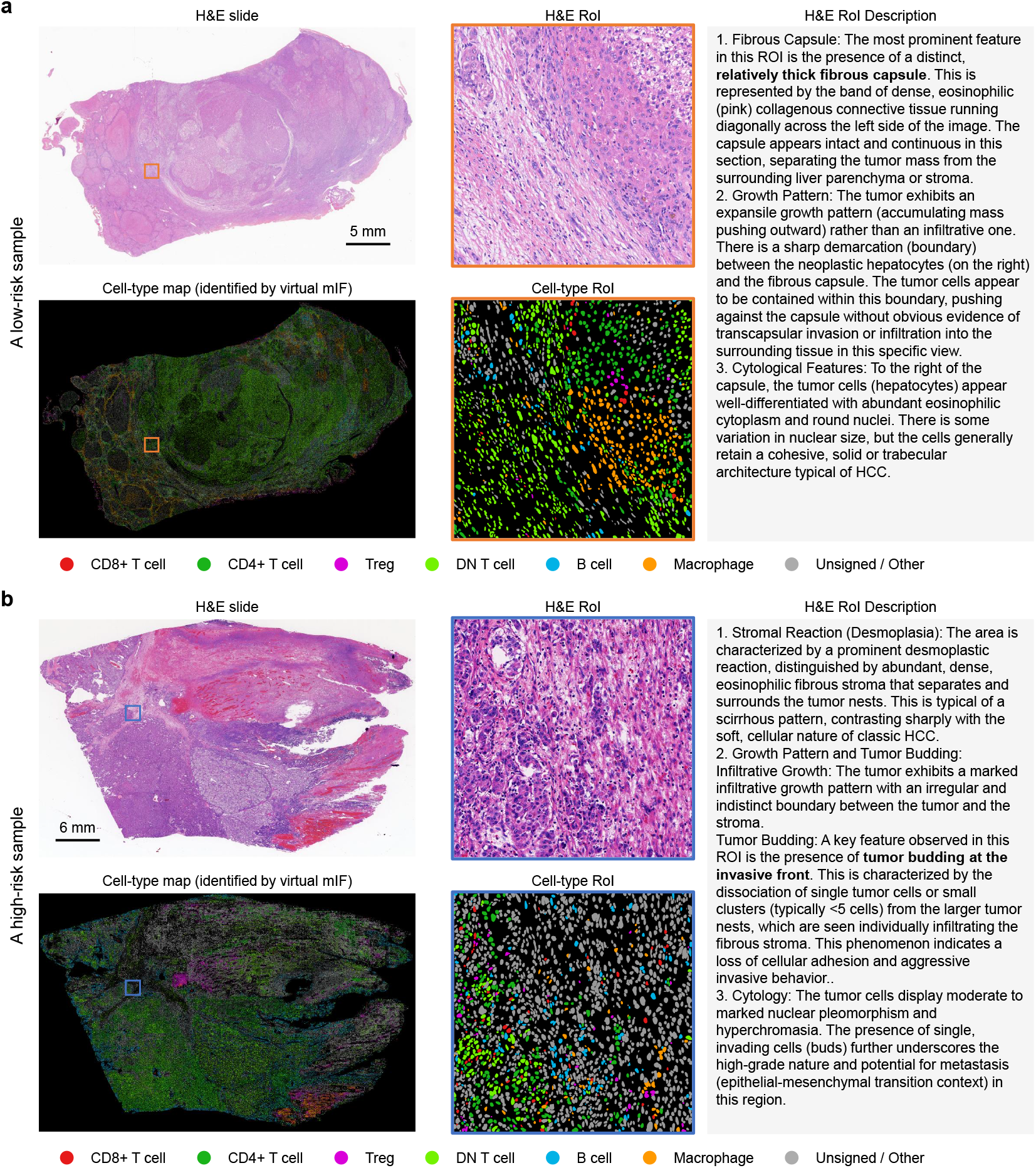
Distinct histological and immune landscapes: Encapsulated containment versus infiltrative tumor budding. a Representative images of a low-risk sample characterized by a desmoplastic fibrous capsule. The H&E RoI (top) reveals an expansile growth pattern where a dense fibrotic rim physically separates the tumor mass from the surrounding liver parenchyma, effectively “walling off” the tumor. The corresponding virtual multiplex immunofluorescence (mIF) map (bottom) uncovers the cellular basis of this “containment niche,” showing a dense infiltration of macrophages and CD4+ T cells co-localized within the fibrotic capsule. **b** Representative images of a high-risk sample exhibiting an aggressive infiltrative and budding pattern. In sharp contrast to the low-risk phenotype, the H&E RoI (top) displays tumor budding at the invasive front, characterized by the dissociation of single tumor cells or small clusters infiltrating the fibrous stroma. The corresponding mIF map (bottom) illustrates a macrophage-depleted microenvironment with a disrupted immune architecture.

**Extended Data Table 1.** Prognostic performance of macrophage abundance (count of macrophage per patch) across four independent cohorts. High-risk and Low-risk mean the number of high-risk and low-risk patients.

| Center | Cutoff | High risk | Low risk | Events | Number of Patients | *P* value | C-index |
| --- | --- | --- | --- | --- | --- | --- | --- |
| FAZJU* | 3.77 | 55 | 55 | 65 | 110 | 0.0000449 | 0.6533 |
| SAZJU | 3.77 | 106 | 105 | 52 | 211 | 0.0447 | 0.5570 |
| TCGA | 3.77 | 266 | 30 | 109 | 296 | 0.0182 | 0.5872 |
| YWCH | 3.77 | 21 | 34 | 35 | 55 | 0.000709 | 0.6155 |

